# Mutations in DNA polymerase δ subunit 1 mediate CMD2-type resistance to Cassava Mosaic Geminiviruses

**DOI:** 10.1101/2022.04.13.487913

**Authors:** Y.W. Lim, B.N. Mansfeld, P. Schläpfer, K.B. Gilbert, N.N. Narayanan, W. Qi, Q. Wang, Z. Zhong, A. Boyher, J. Gehan, G. Beyene, Z.D. Lin, W. Esuma, S. Feng, C. Chanez, N. Eggenberger, G. Adiga, T. Alicai, S.E. Jacobsen, N.J Taylor, W. Gruissem, R.S. Bart

## Abstract

Cassava mosaic disease suppresses cassava yields across the tropics. The dominant *CMD2* locus confers resistance to the cassava mosaic geminiviruses. It has been reported that CMD2-type landraces lose resistance after regeneration through *de novo* morphogenesis. As full genome bisulfite sequencing failed to uncover an epigenetic mechanism for loss of resistance, we performed whole genome sequencing and genetic variant analysis and fine-mapped the CMD2 locus to a 190 kilobase interval. Data suggest that CMD2-type resistance is caused by a nonsynonymous, single nucleotide polymorphism in *DNA polymerase δ subunit 1* (*MePOLD1*) located within this region. Virus-induced gene silencing of *MePOLD1* in a Cassava mosaic disease-susceptible cassava variety produced a recovery phenotype typical of CMD2-type resistance. Analysis of other CMD2-type cassava varieties identified additional resistance alleles within *MePOLD1. MePOLD1* resistance alleles represent important genetic resources for resistance breeding or genome editing, and elucidating mechanisms of resistance to geminiviruses.

## INTRODUCTION

Cassava (*Manihot esculenta* Crantz) is a highly heterozygous staple root crop that feeds nearly a billion people worldwide^1^. Cassava yields are suppressed by infections with cassava mosaic geminiviruses (CMG, Family *Geminiviridae*: Genus *Begomovirus*) which collectively cause cassava mosaic disease (CMD). Eleven species of CMG are known to infect cassava across sub-Saharan Africa, the Indian subcontinent, and recently in several countries of South-East Asia^2^. CMGs possess two circular single-stranded DNA genomes that are transmitted by the whitefly *Bemisia tabaci* and spread by farmers who plant infected stem cuttings to establish the next cropping cycle^3,4^.

Understanding genetic sources for resistance to geminiviruses is critical to securing yields for cassava farmers. Three types of resistance to CMGs have been described in cassava as CMD1, CMD2, and CMD3^5,6^. In all cases the genes responsible for resistance and their modes of action remain unknown. CMD2-associated resistance, which was discovered in landraces collected across West Africa, is a dominant single genetic locus located on Chromosome 12^7–10^. We reported previously that CMD2-type resistance is lost when plants are regenerated through *de novo* morphogenesis in tissue culture^11^ (Fig. 1a). While loss of CMD2 resistance (LCR) occurs consistently in this manner in multiple landraces, LCR was not observed in varieties developed through breeding programs^12^. Epigenetic somaclonal variation is well known to produce phenotypic changes in plants regenerated from *in vitro* cultures^13,14^. We hypothesised, therefore, that the LCR phenotype is caused by culture-induced epigenetic changes at the CMD2 locus. Single-cytosine resolution epigenome-wide association studies (EWAS) were performed on multiple cassava plant lines, before and after *in vitro* morphogenesis. While methylation changes were found across the genome, no consistent methylation changes were observed within the CMD2 locus (Supplementary Fig. 1, Supplementary Table 1).

**Fig. 1.**
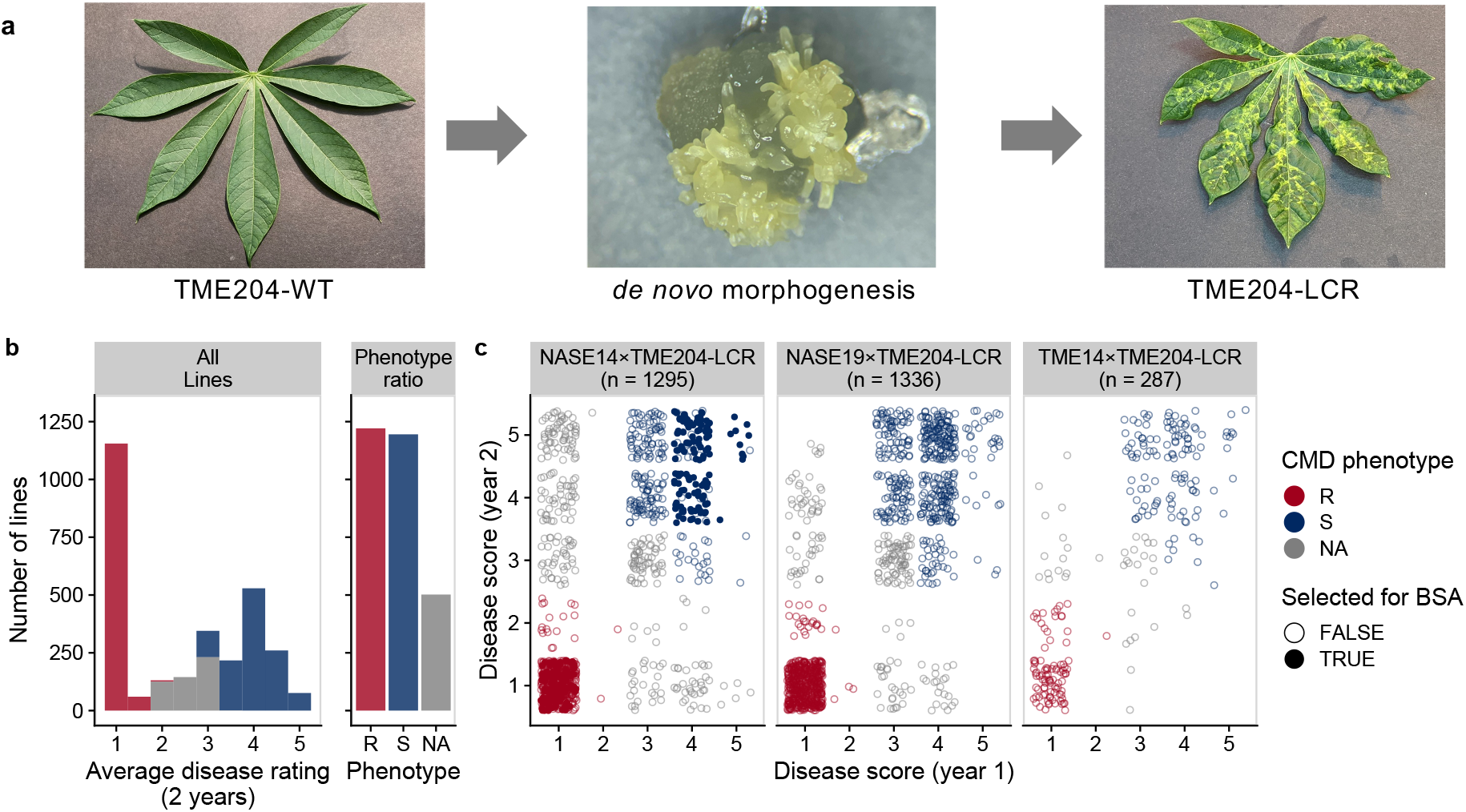
CMD2 type cassava varieties lose resistance upon *de novo* morphogenesis. A) Left – TME204-WT CMD2-type plants challenged with cassava mosaic geminivirus remains symptom free. Middle – embryogenic structures arise from tissue culture induced de novo morphogenesis. Right – Regenerated plant shows classic mosaic symptoms after virus challenge. B) F1 populations derived from heterozygous resistant parents (NASE14, NASE19, TME14) crossed with susceptible loss-of-CMD2-resistance (LCR) line. Plants were grown and phenotyped in the field in Uganda and scored for disease over two years on a 1-5 disease score. The disease rating distribution across all populations segregates at 1:1. !^2^ = 0.59 (C) In each population, ∼15% of lines with consistent phenotypes over the two years were selected for bulk segregant analysis (BSA) mapping (solid circles).

We therefore investigated the relationship between the CMD2 and LCR phenotypes by generating three large mapping populations derived from tissue culture regenerated, CMD susceptible plants (TME204-LCR) crossed with resistant varieties heterozygous for CMD2 (NASE14, NASE19, TME14^8,15^). Field phenotyping was performed over two years at a high CMD pressure location in Uganda, and progeny lines assessed for resistance or susceptibility to CMD (Fig. 1b, Supplementary Data 1). Resistance segregated at 1:1 ratio (Fig. 1b, across all populations, *x*^2^ p-value = 0.59), indicating that the dominant wildtype allele of CMD2 is sufficient to restore resistance, and that the CMD2 and LCR phenotypes are caused by a single genetic locus. If LCR results from a somaclonal epiallele, then passage of CMD resistant F^1^ progeny through morphogenesis would result in the LCR phenotype. However, three independent, resistant F_1_ progeny retained resistance through three consecutive cycles of somatic embryogenesis and plant regeneration, indicating that sexual propagation prevents LCR from occurring after *de novo* morphogenesis in tissue culture (Supplementary Fig. 2). These results indicate that the CMD2 and LCR traits have a genomic basis. We postulate that spontaneous mutation(s) causing CMD2 resistance occurred in the meristems of field grown West African landraces and became fixed as periclinal chimeras (Supplementary Fig. 3). The subset of mutated cells continued to develop into resistant branches that were then selected by farmers and maintained through clonal propagation, as is common in other crop species^16–18^. Loss of resistance to CMD would be explained if *de novo* morphogenesis occurs from cell layers that do not carry the resistance allele. Gametes are typically derived from cells within the L2 layer of the meristem^19^, thus if L2 cells carried the dominant *CMD2* mutation it would be transmitted to the next generation in a Mendelian manner. Resulting progeny plants would not be chimeric for the resistance allele and, as we report here, would not lose resistance to CMD after morphogenesis (Supplementary Fig. 3).

We combined whole genome sequencing and genetic variant analysis (WGS-GVA) with fine-mapping to identify *CMD2* and further understand the LCR trait. WGS-GVA has been used to understand the genetics behind rare human diseases. Causal variants shared by multiple individuals or families are revealed by comparison of WGS from sick and healthy individuals^20,21^. We performed WGS-GVA to identify genetic changes in three CMD resistant and five susceptible F_1_ plants (Supplementary Data 2). A filtering approach (Methods, Expanded Methods, SNP analysis) identified 405 SNPs segregating with the resistance phenotype in these individuals (Supplementary Data 3). We hypothesised that if the LCR phenotype is indeed caused by a mutation within *CMD2*, then susceptible LCR lines should share variants with susceptible F_1_ individuals, while wildtype resistant TME204 would not. Of the 405 SNPs identified in the resistant F_1_ progeny, only one nonsynonymous SNP is heterozygous in the genome of resistant TME204 and absent in the genome of susceptible TME204-LCR plants. This observation is consistent with the hypothesis that CMD resistance is a chimeric trait in landraces and that passage through culture-induced embryogenesis leads to loss of chimerism. The SNP is located in the coding sequence of *MePOLD1* (Manes.12G077400) and changes valine to leucine (V528L) (Fig. 2a). EWAS confirmed that *MePOLD1* has no DNA methylation differences in resistant and susceptible genotypes (Supplementary Fig. 1d).

**Fig. 2.**
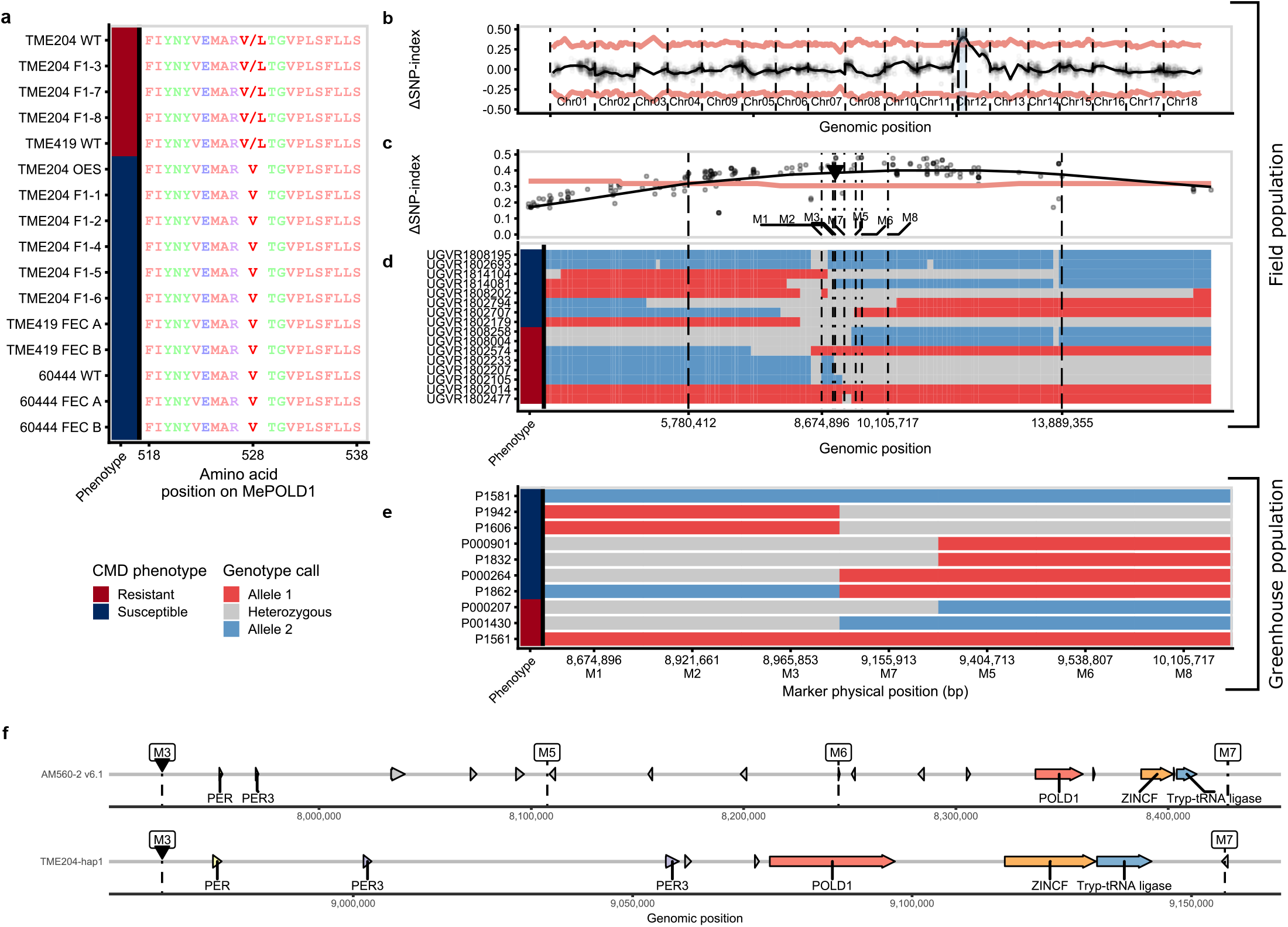
Whole genome sequencing and genome variant analysis (WGS-GVA) and fine mapping reveal nonsynonymous SNPs in MePOLD1 that segregate with resistance. A) TME204-WT and F1 progeny, TME419 WT, 60444 WT and TME204, TME419 and 60444 plants regenerated from tissue culture were tested for resistance and susceptibility. TME204 WT, F1-3, F1-7, F1-8, and TME419-WT plants had CMD2 resistance while all other plants were susceptible to ACMV infections. The resistance phenotype is indicated on the left bar (Red – Resistant; Blue – Susceptible). A haplo-type-restricted G to C transversion in the MePOLD1 gene at location 9,081,215 bp causes a heterozygous V528L mutation in MePOLD1. Two large (n::1,000) F1 mapping populations derived from NASE14xTME204-LCR were used to fine map CMD2 (B-E). B) An *in silico* bulk segregant approach was performed using the field phenotyping and genotyping by sequencing (GBS) data (Fig. 1c). The tricube-smoothed allele frequency enrichment (LiSNP-index) across the TME204 hap1 assembly. In C and D the red line denotes the 95% confidence interval. The highlighted region on Chr12 defines the significantly linked CMD2 region. C) Enlargement of the CMD2 locus mapping results. Each point represents a SNP and its corresponding LiSNP-index. The dashed lines indicate the borders of the mapped locus between ∼5-13Mb. The previously reported associated marker from Rabbi et al., 2020 is indicated by black arrow. D) Examining the GBS SNP data from individual recombinants within the locus improves the mapping resolution to ∼300 kb. Genotypes are extended downstream until the next SNP called. Two non-recombinant homozygous resistant and susceptible lines are added as a control (top and bottom). Based on the location of the mapped locus, and the previously identified GWAS marker, KASP markers (M1-8) were developed for fine mapping (positions denoted by dot-dash lines in C and D). E) A second fine mapping population was phenotyped in the greenhouse using a virus induced gene silencing-based infection assay. Recombinants within the region place CMD2 in the 190Kb interval between markers M3 and M7. Lines P1581 and P1561 are non-recombinant susceptible and resistant controls, respectively. In C and E the genotype at each SNP or marker is indicated by the color (Allele 1, Red, linked to Resistance; Allele 2, Blue, linked to Susceptibility). The resistance phenotype is indicated on the left bar as above. F) Genomic rearrangements within the fine mapped CMD2 locus introduce new gene candidates.

We also pursued fine-mapping to pinpoint the CMD2/LCR genomic location. The recently released haplotype resolved genome assemblies of CMD2-resistant African cultivars TME7^22^ and TME204^23^ were leveraged to perform *in silico* bulk segregant analysis (BSA) (based on Takagi et al (2013)^24^ and Mansfeld and Grumet (2018)^25^) to map CMD2 resistance. First, F_1_ progeny were screened in the field in Uganda and genotyped with GBS (Fig. 1b, Supplementary Data 1). These data co-localize the CMD2/LCR locus with the previously identified CMD2 locus^9^, placing it on Chromosome12 between 5 and 13 Mb of the TME204 haplotype 1 assembly^23^ (Fig. 2b). We identified recombinants within this region using SNP calls from individual samples, thus narrowing the CMD2/LCR-locus to roughly 300 kb (Fig. 2c, d). To more accurately fine-map the locus, kompetitive allele specific PCR (KASP) markers were developed bracketing this region (Fig. 2c-f, Supplementary Fig. 4, and Supplementary Data 4). Approximately 1,000 F_1_ individuals derived from a NASE14×TME204-LCR cross were genotyped and then phenotyped in the greenhouse (Supplementary Data 5) using a previously described virus induced gene silencing (VIGS)-based infection assay^26^. We identified 64 (∼6.57 cM) recombinants between markers M1 and M8 and further screened those individuals using three additional markers (M3, M5, M7). This allowed the identification of recombinants which narrowed the CMD2/LCR locus to 190 kb, between M3 (8,965,853 bp) and M7 (9,155,913 bp) in the TME204-hap1 assembly^23^ (Fig. 2e,f).

The marker order in both TME7 and TME204^22,23^ assemblies is different than in the AM560-2 v6.1 assembly^27^, suggesting a translocation or assembly error in the region which may have complicated previous efforts to find *CMD2* (Fig. 2f). The newly defined fine-mapped locus consists of eight annotated genes, including several peroxidase genes that were previously proposed as CMD2 candidate genes^9,10,28^ and *MePOLD1* (Fig. 2f). Differential gene expression analyses between susceptible and resistant individuals revealed no significant differences for genes found within this region (Supplementary Fig. 5). Nucleotide level comparison of WGS data revealed that the V528L SNP in *MePOLD1* was the only genetic change between these recombinant lines.

Taken together, these data suggest that variation within the *MePOLD1* CDS underlie CMD2-type resistance. Finding a nonsynonymous SNP by WGS-GVA in the precisely mapped CMD2 locus by chance is statistically improbable (P = 6.1×10^−4^, Monte Carlo simulation, n = 100,000). Components of the DNA polymerase complex have been reported previously as required for susceptibility to geminiviruses^29–33^. To understand if this holds true for cassava, we targeted *MePOLD1* for downregulation in the CMD-susceptible cassava variety 60444 using VIGS (*MePOLD1*-VIGS)^34^. After inoculation with *MePOLD1*-VIGS, only 25% (*n* = 40) of 60444 plants showed symptoms of infection compared to plants infected with *GUS*-VIGS (76.7%, *n* = 30) and *African cassava mosaic virus* (ACMV) (100%, *n* = 15). CMD symptom severity after *MePOLD1*-VIGS was also reduced in infected plants of 60444 (Hypergeometric Test, P < 0.05, *n* = 40, Fig. 3 a,b) and virus titre was significantly lower when compared to plants inoculated with control VIGS constructs or unmodified ACMV (Fig. 3c). Importantly, plants of 60444 that displayed CMD symptoms after inoculation with *MePOLD1*-VIGS underwent a recovery phenotype typical of CMD2 resistance and atypical for this highly CMD-susceptible variety (Fig. 3d). While the phenotypic result of *MePOLD1-*VIGS was clear, we did not observe a significant downregulation of *MePOLD1* mRNA levels in 60444 inoculated with *MePOLD1*-VIGS vectors (Supplementary Fig. 6). This may be because *MePOLD1* is already expressed at very low levels in leaf tissues (Supplementary Fig. 7^35^), or reflect inherent complexity associated with using a viral vector to down-regulate a gene required for virus replication (Supplementary Fig. 8). A significant reduction in *Tomato yellow leaf curl virus* (TYLCV) accumulation in *Nicotiana benthamiana* was also observed after POLD was downregulated by *Tobacco rattle virus* (TRV)-mediated VIGS^31^. Since TRV is an RNA virus that replicates via a double-strand RNA intermediate, downregulating POLD with TRV-VIGS will not reduce VIGS-mediated siRNA production because TRV is not dependent on POLD for its replication. Together, our results demonstrate that *MePOLD1*-VIGS is sufficient to provide CMD resistance, although further work is necessary to understand why a RNAi-mediated downregulation of *MePOLD1* expression was not observed.

**Fig. 3.**
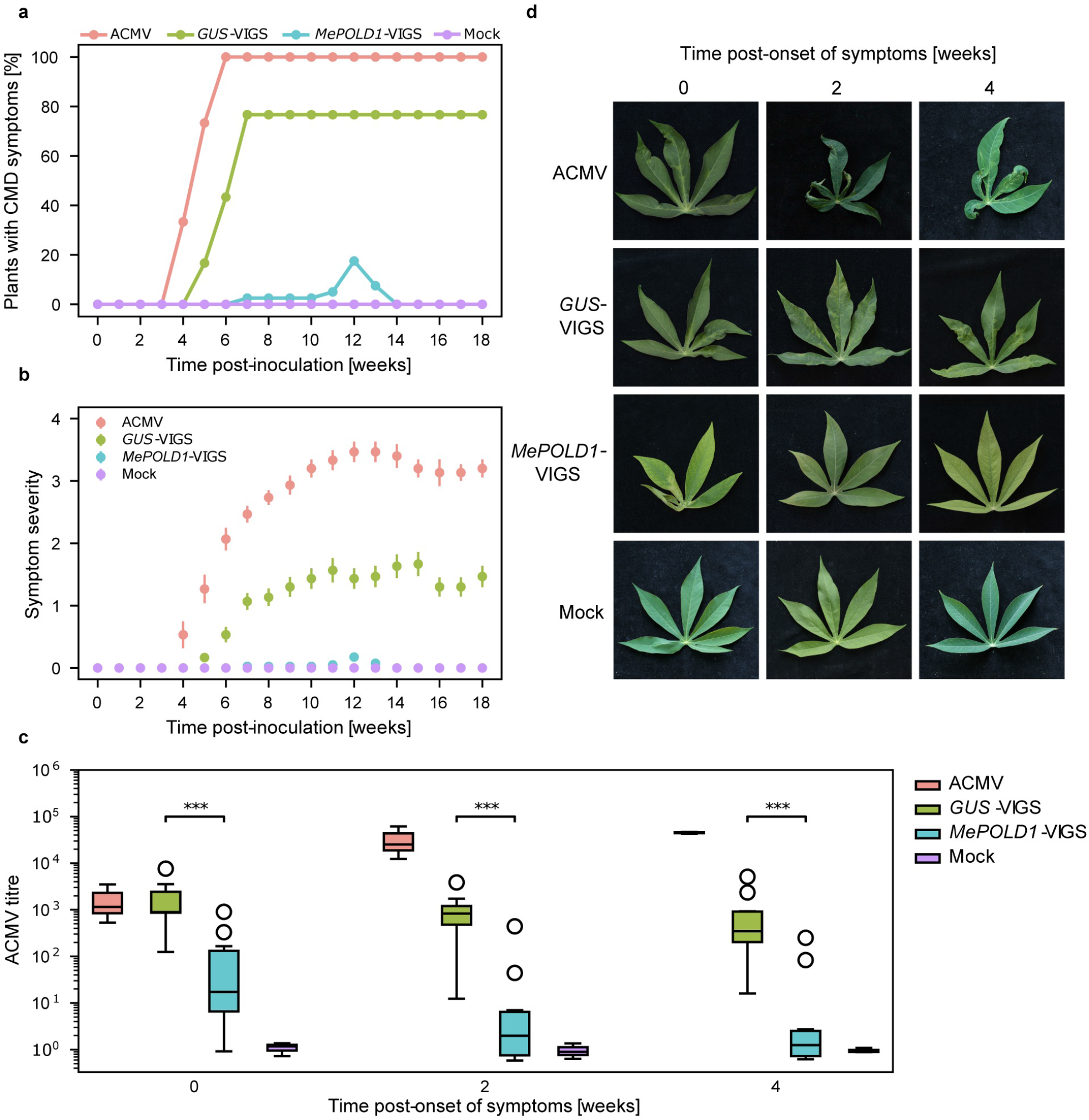
VIGS silencing of *MePOLD1*. CMD-susceptible cassava 60444 recovers from ACMV infection when *MePOLD1* is downregulated by VIGS. (a) Percentage of symptomatic 60444 plants and (b) CMD symptom severity (Fauquet and Fargette, 1990) 18 weeks post-inoculation: ACMV (n = 15), *GUS*-VIGS (n = 30), *MePOLD1*-VIGS (n = 40), and Mock (n = 15). Bars show standard error. (c) Quantification of ACMV titre post-onset of CMD symptoms after inoculation with ACMV (n = 3), *GUS*-VIGS (n = 10), *MePOLD1*-VIGS (n = 10), and Mock (n = 3) (Mann-Whitney U test, ^*^; ^* *^ = P <0.001). Week 0 is the first onset of symptoms detected on individual plants. (d) CMD symptoms on cassava leaves after ACMV-VIGS inoculation of 60444 plants with week 0 being when first symptoms were detected on individual plants.

We next investigated the *MePOLD1* coding sequence of additional CMD-resistant cultivars using WGS-GVA and/or Sanger sequencing (Fig. 4, Supplementary Data 6). The V528L allele present in TME204 was also observed in TME419 (Fig. 4), consistent with these landraces being closely related, and both collected from farmers’ fields in Togo/Benin^3636^. While other resistant varieties did not contain the V528L allele, two additional nonsynonymous SNPs were identified within *MePOLD1* (G680V in TME3, TME8, TME14, NASE12 and NASE14 and L685F in TMS-9102324) (Fig. 4, Supplementary Fig. 9). These results suggest that several distinct *MePOLD1* alleles may explain CMD2 resistance. We also queried publicly available re-sequencing data of diverse cassava germplasm^27,37^ and cross referenced these varieties for CMD severity phenotype data available at CassavaBase^38^. Of the 241 accessions with re-sequencing data, 153 have associated CMD susceptibility scores. *MePOLD1* SNPs were identified in 94 of the resistant accessions (CMD score of less than 2 out of 5). Specifically, 6, 52, and 36 accessions harbour V528L, G680V, or L685F, respectively. (Fig. 4b). Analysis of the remaining 59 varieties identified three additional nonsynonymous SNPs in *MePOLD1*, unique to accessions with CMD severity scores below 2: L598W, G680R, and A684G; found in 17, 2, and 4 samples, respectively (Fig. 4c). In every case, across 117 samples in which POLD1 variants were identified, the putative resistance allele is observed in the heterozygous context, suggesting that these amino acid changes might be deleterious if homozygous. Indeed, an EMS mutant in Arabidopsis POLD1 (at position A684 in MePOLD1; Fig. 4c) is hypomorphic and lethal at 28oC^39^. Five of the six mutations identified in MePOLD1 (V528L, G680V, G680R, A684G, L685F) are immediately adjacent to the R696-E539 (MePOLD1: R681-E524) salt bridge between the finger and N-terminal domains described in yeast POLD (Fig. 4d, e). Mutations disrupting this salt bridge have been shown to result in decreased polymerase activity and fidelity^40,41^. Furthermore, a homozygous R696W mutation is lethal in yeast and is associated with oncogenesis in humans^41^.

**Fig. 4.**
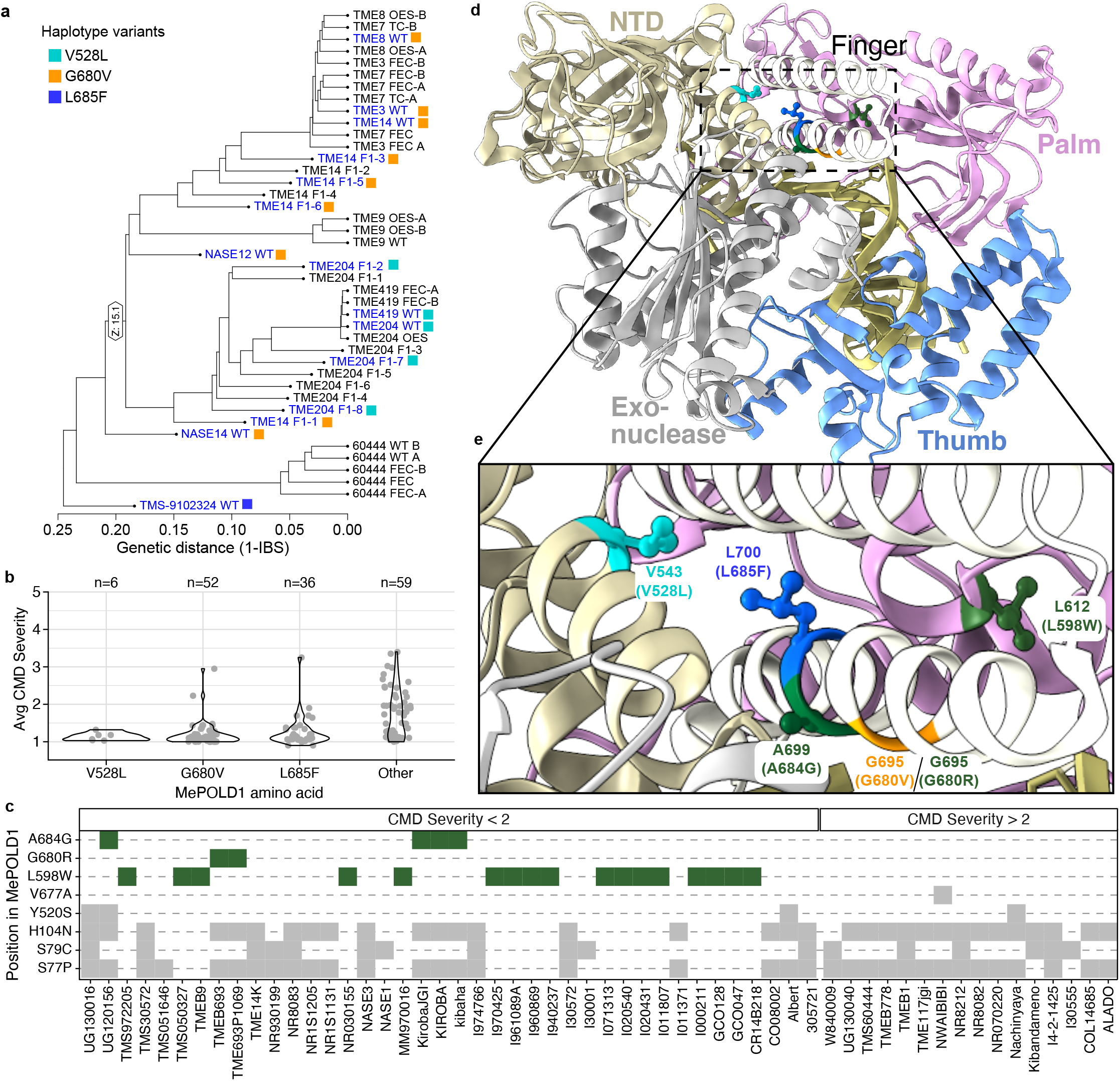
Nonsynonymous SNPs in MePOLD1. (a) Dendrogram of *Manihot esculenta* cultivars analyzed by whole genome sequencing. Non-synonymous SNPs (nsSNPs) in MePOLD1 of various cultivars segregate with CMD2 resistance. Names of resistant cultivars are in blue and harbor either the V528L (cyan), G680V (orange), or L685F (blue) mutation. (b) Average CMD severity across a diverse set of cassava cultivars from the HapMapII population (Ramu *et al*., 2017) that have either one of the three mutations from (a) or an unknown nsSNP in MePOLD1 (“Other”). (c) Identity of all nsSNPs in MePOLD1 of varieties from the “Other” category in (b). Varieties are split by CMD severity score, where less than 2 and above 2 are resistant and susceptible, respectively. In green are the nsSNPs found only in cultivars with CMD severity scores below 2; all other nsSNPs are in gray. (d) Three-dimensional structure of *S. cerevisiae* POLD1 (PDB: 3IAY) with corresponding MePOLD1 mutations highlighted; V528L in cyan, G680V in orange, and L685F in blue. Additional residues identified in (c), L685F and L598W, are in green. Residue name and position in ScPOLD1 are noted and the corresponding information for MePOLD1 is in parentheses. POLD1 functional domains, N-terminal (beige), exonuclease (grey), and structural motifs of the polymerase domain, palm (pink), fingers (white), and thumb (blue), are highlighted. (e) Zoomed in view of the 3D structure centered on the mutated residues found in MePOLD1.

The above data suggest a model wherein *MePOLD1* is a susceptibility factor involved in cassava geminivirus replication and that nonsynonymous mutations within *MePOLD1* lead to CMD2-type resistance. We applied this model to an unexplained observation. The resistant NASE14 parent from the mapping populations is heterozygous for the G680V mutation. NASE14 (the line formerly known as MM96/4271) was developed by crossing in a breeding program at the International Institute for Tropical Agriculture^42^ and does not lose resistance after passage through culture-induced morphogenesis^12^. However, we previously reported an exception: in experiments where NASE14 was used to generate transgenic lines all but one of the transgenic events remained resistant to CMD^43^. To understand this unexpected outcome, targeted Sanger sequencing of *MePolD1* was performed on the transgenic line 5001-NASE14-#41 that had lost CMD2 resistance. The result confirmed that this line retained the heterozygous G680V mutation characteristic of the resistant NASE14 cultivar. However, examining the cloned, full length CDS revealed the presence of an additional heterozygous SNP not present in WT NASE14. This new SNP introduces a premature stop codon at amino acid position 574 within the resistance allele (Supplementary Fig. 10). Thus, transgenic event 5001-NASE14-#41 contains a susceptible version of *MePOLD1*, but lacks its original functional resistance allele, which would explain its acquired susceptibility to infection by CMGs. This spontaneous knock-out of the resistance allele provides further strong evidence that mutations in *MePOLD1* explain CMD2 type resistance in cassava.

Collectively, our data indicate that amino acid changes near the active centre of MePOLD1 cause the dominant CMD2-type resistance. Several dominant resistance genes for plant viruses have been reported, most of which belong to the NBS-LRR class of proteins^44^. MePOLD1 represents an unexpected, novel type of resistance protein in plants. Evidence suggests that this has been selected as a chimeric clonal variant multiple times by West African farmers, and due to its monogenic, dominant nature is now favoured in breeding programs in Africa, India, and South-East Asia^8^. Mutations in POLD predispose humans and mice to a range of cancers, especially mutations that specifically affect the proofreading activity or dNTP selectivity of the enzyme^45^. It is possible that the identified mutations in *MePOLD1* may similarly introduce replication errors in the geminiviruses, which would impair their replication efficacy and thereby reduce virus load in the host plant. This hypothesis is supported by the co-localization of MePOLD1 mutations to those in yeast and humans known to decrease DNA replication activity, and accuracy^40,41,45^. We cannot exclude, however, that the *MePOLD1* mutations weaken or block interactions with the virus replication-enhancer protein AC3, which interacts with subunits of POLD^31^. CMD2 resistance has remained robust in farmers’ fields over at least two decades. However, some caution for overreliance on CMD2 is presented here with evidence that yields and livelihoods for millions of cassava farmers are being secured by a few SNPs in one gene. The identification of mutations in *MePOLD1* as the cause for CMD2-type resistance will facilitate the production of CMD resistant cassava varieties by SNP-assisted breeding or genome editing to introduce the identified SNPs into susceptible cultivars and provides opportunity to further elucidate mechanisms of resistance to geminiviruses.

## METHODS (see Supplementary File 1 for expanded methods section)

### Plant lines, mapping populations and disease scoring

For detailed descriptions of each plant line used in this study, see Supplementary Table 1 and Supplementary Data 1 and 2. TME204-LCR was described previously^46^.

A crossing program was conducted in Uganda during the 2017/2018 cropping season to perform controlled crosses between CMD susceptible cultivar TME204-LCR and three CMD resistant wildtype cassava varieties (TME14, NASE14, NASE19) following the standard procedures described by Kawano (1980)^47^ and Hahn et al (1980)^48^. During the pollination period, special care was taken to cover mature flowers with pollination bags 2-3 days before and after pollination. A total of 7,200 botanical seeds were harvested from mature fruits within three months after pollination and stored in paper bags for approximately three weeks to break dormancy. All seeds were planted in field-conditioned nursery beds and 4,300 resultant seedlings transplanted to a field at six weeks or age and allowed to grow under natural field conditions for 12 months. The field trials were conducted at Namulonge, central Uganda, which is a hotspot for cassava mosaic disease with high whitefly vector populations. CMD-symptomatic plants of local cultivar Bao were planted as spreader rows to augment field inoculation of CMGs. To achieve phenotyping, monthly CMD severity was scores (starting from 1 month after transplanting seedlings) were recorded on a 1-5 scale^49^ where: 1 = no symptoms; 2 = mild chlorotic pattern over the entire leaf although the leaf appears green and healthy; 3 = moderate mosaic pattern throughout the leaf, narrowing and distortion in the lower one-third of leaflets; 4 = severe mosaic, distortion in two-thirds of the leaflets and general reduction in leaf size; and 5 = severe mosaic distortion in the entire leaf. The final CMD severity data recorded at the crop age of 11 months were used for subsequent analyses.

A similar crossing program was established at Kandara, Kenya in which TME204-LCR was crossed with the two CMD resistant wildtype cassava varieties (TME14 and NASE14). Resulting seeds were collected and shipped to DDPSC, St Louis, USA.

### Epigenome-Wide Association Studies (EWAS)

Whole genome methylation of TME7 and TME204 background samples were prepared with Bisulfite Kit (Qiagen, Germantown, Maryland, USA) and enzymatic Methyl-Seq kit (New England BioLabs, Ipswich, Massachusetts, USA), respectively. For more information on library preparation see expanded methods (Supplementary File 1). DNA methylation level at each cytosine was calculated by number of methylated C vs. total C and T count. Differentially Methylated Cytosines (DMCs) were identified by methdiff.py in BSMAP^50^ where differences in CG, CHG, and CHH methylation were at least 0.3, 0.2, and 0.1, respectively. Methylation levels of DMCs of each sample versus three TME7 and one TME204 wildtype were merged as a consensus DMCs table. Methylation levels of each sample in DMCs table were subjected to one-way ANOVA test by comparing seven resistant vs. seven susceptible samples to calculate p-value of each DMC. Manhattan plot of p-value were generated by R package qqman^51^. Methylation track files were visualised with Integrative Genomics Viewer (IGV, v3.0)^52^.

### CMD resistance across cycles of somatic embryogenesis

The three CMD resistant F1 progeny lines, NASE14×TME204-LCR.82, NASE14×TME204-LCR.73 and NASE14×TME204-LCR.16 were established, and micro propagated in tissue culture. Organised somatic embryos (OES) were induced from leaf explants and plants regenerated to produce Cycle 1 OES-derived plants^53^. This process was repeated with Cycle 1 OES plants to produce Cycle 2 OES plants, and again to generate Cycle 3 OES plants for each of the F1 progeny lines. Regenerated plants were established in the greenhouse^53^ and inoculated with *East African cassava mosaic virus* (EACMV-KE2) isolate K201 as described previously^26^. Ten plants were inoculated from each cycle of OES-derived plants for all three progeny and assessed for development of CMD leaf symptoms over a period of 90 days using a 0-5 visual scoring method^54^. At 51 days after inoculation plants were ratooned (cut back) and a new round of CMD symptoms scored on leaves produced by shoot regrowth to confirm the original phenotype.

### Whole genome sequencing and genomic variant analysis

Illumina sequencing: Leaf material was collected from 42 cassava genotypes and FEC material from two cassava genotypes (Supplementary Data 2) for whole genome Illumina sequencing (see Expanded Methods). DNA libraries were prepared using the Illumina TruSeq Nano DNA High Throughput Library Prep Kit (20015965, Illumina, San Diego, California, USA). Libraries were sequenced using an Illumina NovaSeq system for 2 × 151 cycles. On average 100X Illumina paired-end (PE) data were collected per sample.

Pre-processing and mapping of reads was performed using ezRun (https://github.com/uzh/ezRun) in combination with SUSHI^55^. Technical quality was evaluated using FastQC (v0.11.7). Possible contaminations were screened using FastqScreen (v0.11.1) against customised databases (See Expanded Methods). Reads were pre-processed using fastp (v0.20.0) and aligned to the *Manihot esculenta* TME204 genome (V1.0, FGCZ) using Bowtie2 (v2.3.2) with the “--very-sensitive” option. PCR-duplicates were marked using Picard (v2.9.0). Frequency-based calls for all variants with allele frequency above 20% were performed with freebayes-parallel (v1.2.0-4-gd15209e). Relatedness analysis of SNPs using identity-by-descent (IBD) measures, was performed using the R/Bioconductor Package SNPRelate (v 3.13).

SNP analysis: To find potential SNPs, a custom python script (https://github.com/pascalschlaepferprivate/filter_vcf) parses the VCF file produced by freebayes, computes total coverage of the SNP, and then absolute and relative read coverage of all SNP variants. Samples were organized as ingroup (genotypes that show a SNP variant of interest), outgroup (genotypes that do not show SNP variant of interest), facultative ingroup (genotypes that may show SNP variant of interest), and facultative outgroup (genotypes that may not show SNP variant of interest), and SNPs were filtered according to these groups and additional parameters (see expanded methods in Supplementary File 1).

### Genetic mapping

Genotyping by Sequencing and *in silico* bulk segregant analysis: Approximately 1,300 individual F_1_ progeny and the parental lines from the NASE14xTME204-LCR population generated in Kenya were characterised with genotyping-by-sequencing (GBS) at UW-Madison Biotechnology Center following their standard ApeKI restriction enzyme protocol. Reads of 100bp were demultiplexed^56^ and mapped^57^ to the TME204 hap1 assembly^23^. SNPs were called using GATK4^58,59^ and quality filtered SNPs that were heterozygous in both parents retained using vcftools v0.1.14^60^. Using the field phenotypes, a random subset of the most CMD resistant and most susceptible lines was selected as the resistant and susceptible bulks (n = 125 each), respectively, to perform *in silico* bulk segregant analysis using the QTLseqr package^25^.

Fine-mapping using GBS and KASP markers: To further narrow the CMD2 locus, individual F_1_ progeny were analysed for recombination events within the defined locus (∼5-13Mb). While mapping in outcrossers using F_1_ populations is established, mapping in this population is complicated by the TME204-LCR parent in that heterozygous progeny can be either resistant or susceptible. Thus, only recombinants with a genotype-phenotype mismatch were selected as informative. For example, in a phenotypically resistant F_1_ line with a recombination that transitions from genetically heterozygous to genetically homozygous susceptible, one can exclude the homozygous susceptible region as not carrying *CMD2*. Six resistant and six susceptible recombinant individuals were identified with such recombination within the broad CMD2 locus and were used to exclude genomic regions in which at least two lines supported such exclusion. The narrow locus defined by GBS (Chromosome12: 8,976,221-9,314,764) was used to design KASP markers (Supplementary Data 4) spanning 1.5 Mb bracketing this region. Additional recombinants were sought in a similar manner within a second ∼1000 individual population using highly accurate genotyping and phenotyping assays (KASP-marker-based assay combined with phenotyping with a VIGS based approach^26^). All recombinants were sequenced using Illumina WGS data and nucleotide level comparison was performed by alignment to TME7^22^ and TME204^23^ assemblies and manual inspection using CLC Genomics and IGV^52^.

Phenotyping for fine-mapping: F_1_ progeny seeds were germinated in a growth chamber at DDPSC, transferred to the greenhouse and inoculated with a virus-induced-gene-silencing version of *East African cassava mosaic virus* K201 (SPINDLY-VIGS), as described by Beyene et al. (2017). Plants were assessed over a four-week period. Plants which died were scored as CMD susceptible while those that recovered from initial symptoms and re-established healthy growth were scored as CMD resistant.

### RNAseq analysis

For differential expression analysis, first a transcriptome fasta of the spliced exons was made from the TME204-hap1 gff file using ‘gffread -w’ from the cufflinks package^61^. This transcriptome was then concatenated to the whole genome to prepare an alignment decoy file and index using the commands here https://combine-lab.github.io/alevin-tutorial/2019/selective-alignment/. Trimmed RNAseq reads were then pseudo-aligned to the TME204-hap1 transcriptome using Salmon v1.5.2 default settings^62^. Read count data was imported into R using the tximport package^63^. Samples were then defined as resistant or susceptible and differential expression on the integer count values was performed using DESeq2^64^. Genes with a sum of less than 50 reads across all samples were excluded from analysis. Differential expression was performed using “apeglm” as the Log Fold Change Shrinkage method^65^. Genes were defined as being significantly differentially expressed if they had an adjusted p-value^66^ of less than 0.05. Normalised counts were plotted using ggplot and tidyverse^67^ functions in R.

### VIGS targeting of *MePOLD1*

A VIGS approach was designed and performed based on Lentz et al. (2019)^34^. A 400 bp coding sequence of *MePOLD1* (position 438-837, corresponding to 8905774-8905965 of chr12 in AM560 v8, 9076083-9076741 of chr12 in TME204 hap1) as synthesised (Twist Biosciences, California, USA) and inserted in the multiple cloning site of the ACMV-based VIGS vector using KpnI and SpeI. The 400 bp coding sequence is conserved in MePOLD1 of 60444, TME3, TME204 and AM560 and n-mers (18 – 24 nt) were checked against the cassava AM560 v6.1 genome sequence with SGN VIGS from Sol Genomics (^https://vigs.solgenomics.net/)38^ to validate that the sequence we used to target MePOLD1 has no off-targets in cassava. The number of 60444 plants inoculated were *n* = 15 for ACMV, *n* = 40 for *MePOLD1*-VIGS, *n* = 30 for *GUS*-VIGS, and *n* = 15 for Mock treatments. Leaf symptom scoring was based on Fauquet and Fargette (1990)^54^. ACMV titre and *MePOLD1* expression were quantified through qPCR from total DNA and RNA extracted respectively from the top 1-2 leaves harvested at first signs of CMD symptoms. A Mann-Whitney U test was used to analyse the statistical significance. Primers are listed in Supplementary Table 2.

### Identification of additional MePOLD1 variants

A publicly available dataset was accessed containing sequencing data of 241 diverse accessions that identified over 28 million segregating variants^37^. All positions within the *MePOLD1* gene (AM560-2 v6.1 coordinates) were extracted from the Chromosome12 VCF file available through the cassavabase.org FTP server (c12.DepthFilt_phasedSNPs.vcf), and effects of the variants on the protein coding sequence determined using snpEff^68^. Additional analysis was done with Sanger sequencing (Supplementary Data 6). Names listed in Fig. 4c are as listed in Ramu et al.^37^ We note that according to this publication, TMS972205 contains a different SNP than the one identified here and is referred to as TMS-972205.

### POLD1 Protein sequence analyses

The 3D structure of the yeast POLD catalytic subunit and template DNA (PDB ID: 3IAY, Swan et al., 2009), was visualised in ChimeraX^69^. The N-terminal domain, exonuclease domain, and finger, palm, and thumb motifs from Swan et al., 2009^70^ were colour-coded and the residues corresponding to the nonsynonymous mutations identified across the cassava varieties are highlighted.

### Analysis of *MePOLD1* in 5001-NASE 14-#41

The full-length cDNA of *MePOLD1* was amplified from cassava plant line 5001-NASE 14-#41^43^. Primers were designed to be specific for the haplotype carrying the resistance *MePOLD1* allele and PCR performed. The PCR product was cloned into the binary vector pCAMBIA1305.1 using the In-Fusion® HD Cloning Kit (Takara Bio USA, Inc.) and the resulting clones sequenced by Sanger sequencing. Primers are listed in Supplementary Table 2.

## DATA AVAILABILITY

Source data are provided with this paper as Supplementary Datasets. Raw bisulfite sequence data is available through NCBI GEO (GSE192748 data will be made public before publication). Whole Genome Sequencing and RNAseq raw read data can be accessed at NCBI (https://www.ncbi.nlm.nih.gov/bioproject/PRJNA787456;

Reviewer link: https://dataview.ncbi.nlm.nih.gov/object/PRJNA787456?reviewer=r6e2b80b8055v1lcugaa2q51lh).

## CODE AVAILABILITY

Scripts used to generate figures are deposited in github and in the Supplementary Data available with this publication.

## ACKNOWLEDGEMENTS

Identification of the CMD2 resistance genes was supported by a grant from the Bill & Melinda Gates Foundation to ETH Zurich (Investment INV-008213), funding from ETH Zurich and the Donald Danforth Plant Science Center, Institute for International Crop Improvement. W.G. is supported by a Yushan Scholarship of the Ministry of Education in Taiwan. We thank Joel Kuon (ETH Zurich) for initial investigation of the cassava CMD2 locus and Emily McCallum (ETH Zurich) for Sanger sequencing of genes in the CMD2 region, and the high throughput sequencing team at the Functional Genomics Center Zurich for Illumina sequencing. We also thank Irene Zurkirchen (ETH Zurich) for greenhouse propagation and maintenance of the cassava plants, Justin Villmer, Jennifer Winch and Claire Albin (DDPSC) for plant regeneration, propagation, greenhouse care and virus inoculations, and Douglas Miano, Catherine Taracha, Paul Kuria and Theresia Munga (Kenyan Agriculture and Livestock Research Organization, Kenya) for production of F_1_ progeny lines in Kenya.

## AUTHOR CONTRIBUTIONS

Y.W.L. contributed to the WGS-GVA that led to the identification of MePOLD1 resistance alleles, designed the VIGS experiments, analysed data and co-wrote the manuscript. B.M. designed and performed the rough and fine mapping, the transcriptomics, analysed data and co-wrote the manuscript. P.S. conceived, designed, and performed the WGS-GVA that led to the identification of the MePOLD1 resistance alleles, analysed data and co-wrote the manuscript. K.B.G. designed and performed the analysis of publicly available re-sequencing and contributed to writing the manuscript. N.N. designed, performed, and analysed greenhouse CMG experiments and RNAseq datasets. Q.W. performed sanger sequencing, cloned, and sequenced the full length *MePOLD1* cDNA from 5001-Nase14-#41 and contributed to writing the manuscript. Z.Z. performed the EWAS and contributed to writing the manuscript. A.B. developed pipelines for and performed analysis of GBS data, SNP calling and rough mapping. J.G. performed and contributed to the development and analysis of the KASP fine-mapping markers. G.B. designed, performed, and analysed RNAseq, SPINDLY-VIGS experiments and contributed to design of field crossing programs. Z.D. L. performed analysis on all lines with sanger sequencing, analysed data and contributed to writing the manuscript. W.E. co-designed field phenotyping experiments, performed mapping population and genetic crossing field experiments, and analysed data. S.F. constructed and sequenced whole genome bisulfite libraries and assisted in data analysis. W.Q. performed the Illumina read mapping, variant calling and clustering analysis. C.C. prepared the Illumina sequencing and performed VIGS experiments. N.E. performed VIGS experiments. G.A. performed field phenotype data collection from mapping populations, prepared samples for sequencing and contributed to data analysis. T.A. conceived and co-designed field phenotyping experiments, analysed data and contributed to writing the manuscript. S.E.J. conceived and designed the methylation experiments, analysed data, and contributed to writing the paper. N.J.T. conceived and designed plant tissue culture investigations, co-designed field crossing and CMD resistance experiments, analysed data and co-wrote the manuscript. W.G. conceived, designed, and managed the collaborative research project and efforts between groups, contributed to the design of experiments, analysed, and interpreted data, and co-wrote the manuscript. R.S.B. conceived and designed experiments, analysed data, coordinated efforts between groups and co-wrote the manuscript. The authors wish it to be known that Y.W.L., B.M. and P.S. are equal first authors and that W.G. and R.S.B. are equal last and corresponding authors. For the purpose of their CVs, they may list their names as first or last author positions.

