## Supplemental Table 1, 2 and Figures 1-10 for "Mutations in DNA polymerase δ subunit 1 mediate CMD2-type resistance to Cassava Mosaic Geminiviruses"

### Supplementary File Contents:

- Expanded Methods
- Supplementary Table 1
- Supplementary Table 2
- Supplementary Figures 1-10

### Expanded Methods

<u>Plant lines and mapping populations and disease scoring</u>

The disease rating distributions of the entire ~3000 individual population were plotted to assess if epistatic segregation ratios could be observed. To ensure robust resistance phenotype descriptions, only plants with a two-year mean disease rating of less than 2 were defined as resistant and lines with consistent disease ratings above 3 in both years were denoted as susceptible. The 1:1-R:S ratio was tested using a chi-square test (chisq.test function) in R.

<u>DNA methylation library preparation and EWAS analysis</u>

Genomic DNA from samples in the TME7 background were end-repaired and ligated with TruSeq DNA single adapters (Illumina) using a Kapa DNA HyperPrep kit (Roche). Adapter-ligated DNA was converted with an EpiTect Bisulfite Kit (Qiagen). Converted DNA was PCR-amplified by MyTaq polymerase (Bioline) for 12 cycles. EM-seq libraries for samples in TME204 background were prepared from sheared DNA using an enzymatic Methyl-Seq kit following manufacturer instructions (New England BioLabs) with 6 PCR cycles[1]. The libraries were run on D1000 ScreenTape (Agilent) to determine the quality and size, and then purified by AMPure XP beads (Beckman Coulter). Library concentrations were measured with a Qubit dsDNA Broad-Range Assay kit (ThermoFisher). Libraries were sequenced on a HiSeq 2500 or NovaSeq 6000 sequencer (Illumina).

WGBS and EM-seq reads were mapped to haplotype 1 and haplotype 2 genomes of TME204 by BSMAP (v2.90)[2] allowing 0 mismatches and one best hit (-v 0 -w 1)[2]. Duplicated reads were removed with SAMtools (v1.3.1)[3]. Reads with three or more consecutive methylated CHH sites were considered as unconverted reads and removed in the following analysis. Conversion rate was estimated by calculating methylation level of the chloroplast genome.

Whole genome sequencing and genomic variant analysis

Illumina sequencing: Leaf material was collected from 42 cassava genotypes and friable embryogenic callus (FEC) material from two cassava genotypes (Supplementary Table 3) for whole genome Illumina sequencing. DNA was extracted using the DNeasy Plant Mini Kit (QIAGEN, Germany). DNA samples were sent to the Functional Genomics Center Zurich (FGCZ) for Illumina sequencing. DNA libraries were prepared using the Illumina TruSeq Nano DNA High Throughput Library Prep Kit (20015965), following the manufacturer's protocol (Illumina, San Diego, California). Libraries were sequenced using an Illumina NovaSeq system for 2 × 151 cycles, according to the manufacturer's instructions (Illumina, San Diego, California). On average 100X Illumina paired-end (PE) data were collected per sample.

Pre-processing and Mapping of reads: Quality control and Bowtie2 alignment of the Illumina PE reads were performed using data analysis workflows in the R-meta package ezRun (https://github.com/uzh/ezRun), managed by the data analysis framework SUSHI[4], which was developed and maintained by FGCZ. Technical quality was evaluated using FastQC version 0.11.7. Possible contaminations were screened using FastqScreen version 0.11.1 against a customized database in SUSHI, which consists of SILVA rRNA sequences (https://www.arb-silva.de/), UniVec (https://www.ncbi.nlm.nih.gov/tools/vecscreen/univec/), refseq mRNA sequences and selected genome sequences (human, mouse, Arabidopsis, bacteria, virus, phix, lambda, and mycoplasma) (https://www.ncbi.nlm.nih.gov/refseq/). Illumina PE reads were pre-processed using fastp (v0.20.0), where sequencing adapters and low-quality ends (<Q20) were trimmed. Trimmed reads passing the filtering criteria (average quality >= Q20, minimum length >=18 bp) were aligned to the *Manihot esculenta* TME204 genome (V1.0, FGCZ) using Bowtie2 version 2.3.2 with the --very-sensitive option. PCR-duplicates were marked using Picard version 2.9.0. Read alignments were comprehensively evaluated using the mapping QC app in SUSHI, in terms of different aspects of DNA-seq experiments, such as sequence and mapping quality, sequencing depth, coverage uniformity and read distribution over the genome.

Freebayes Variant Calling: Multi-sample, frequency-based calls for all variants with allele frequency above 20% were generated using the freebayes-parallel script in freebayes (v1.2.0-4-gd15209e), with 24 threads of freebayes running in parallel across regions of 100kb in the reference genome. Dendrogram and underlying relatedness analysis of SNPs using identity-by-descent (IBD) measures was performed using the R/Bioconductor Package SNPRelate (v 3.13).

SNP analysis: To find potential SNPs, a custom python script (https://github.com/pascalschlaepferprivate/filter_vcf) parses the VCF file produced by freebayes, computes total coverage of the SNP, and then absolute and relative read coverage of all SNP variants. Four groups of genotypes can be defined to filter SNP results in the VCF file: ingroup (genotypes that show a SNP variant of interest), outgroup (genotypes that do not show SNP variant of interest), facultative ingroup (genotypes that may show SNP variant of interest), and facultative outgroup (genotypes that may not show SNP variant of interest). Seven parameters are given to the script. Minimal total read coverage (mtrc) defines the minimum number of reads (all variants included) that each genotype has to show to be qualified for further filtering. Minimum relative read coverage (mrrc) in ingroups defines the relative number of times that a SNP variant of interest had to be sequenced in ingroup and facultative ingroup respectively. Maximum absolute noise read coverage (mnrc) is the number of times that a SNP variant of interest is allowed to be sequenced in outgroup and facultative outgroup respectively. The four remaining parameters are minimum number of ingroup hits (ni), the number of genotypes in the ingroup that need to show a SNP variant and equivalent parameters for outgroup (no), facultative ingroup (nfi), and facultative outgroup (nfo). Every SNP is evaluated according to the filtering set by the authors. To identify SNP variants of interest using TME204 germplasm, we used TME204 F1-2, -7, and -8 as ingroup, and TME204 F1-1, -3, -4, -5 and -6 as outgroup and left facultative groups blank. Parameters were set to mtrc = 20, mnrc = 2 (10%), mrrc = 0.2, ni = 3, no = 5, nfi = 0 and nfo = 0. To shortcut the parameter settings and produce the results of the manuscript directly, use option -s TME204. To find SNP variants of interest for TME14, we used

TME14 F1-1, -3, -5, and -6 as ingroup, TME14 F1-2, -4 as outgroup and 60444 FEC Plant A, FEC Plant B, TME3 FEC A, FEC B, TME7 FEC, TME7 FEC Plant A, FEC Plant B, TME8 OES Plant A, OES Plant B, TME9 OES Plant A, OES Plant B, TME204 OES Plant, TME204 F1-1, -3, -4, -5, -6, TME419 FEC Plant A, and FEC Plant B. Parameters were set to mtrc = 20, mnrc = 2 (10%), mrrc = 0.2, ni = 4, no = 2, nfi = 0 and nfo = 9. To shortcut: -s TME14. To find the SNPs for TMS-9102324, ingroups were defined to be TMS-9102324 WT respectively. Outgroup was defined to be 60444 WT, TME14 F1-2, -4, TME204 F1-1, -3, -4, -5, and -6. No facultative ingroup was defined and facultative outgroup consisted of 60444 FEC Plant A, FEC Plant B, TME3 FEC Plant A, FEC Plant B, TME7 FEC, FEC Plant A, FEC B, TME8 OES Plant A, OES Plant B, TME9 OES Plant A, OES Plant B, TME204 OES Plant, TME419 FEC Plant A, FEC Plant B. Parameters were set to mtrc = 20, mnrc = 2 (10%), mrrc = 0.2, ni = 1, no = 8, nfi = 0 and nfo = 12. To shortcut: -s 91-02324.

<u>Genetic mapping</u>

Reads were demultiplexed into sample fastq files using GBSX v1.3 [5] and mapped to the TME204 hap1 assembly. The GATK4 best practices [6,7] pipeline was followed with one GBS pertinent modification (alignments were not deduplicated) to call SNPs vs the assembly. Using vcftools v0.1.14[8], the SNPs from the parental lines (NASE14 and two TME204-LCR lines) were extracted from the quality filtered ('QD < 2.0, QUAL < 30.0, SOR > 3.0, FS > 60.0, MQ < 40.0, MQRankSum < -12.5, ReadPosRankSum < -8.0') VCF file and filtered to extract only those which are heterozygous in both parents (i.e. pseudo-testcross). The subset of F1s derived from these parental lines (n = 1,295) was extracted and only the pseudo-testcross positions established above were retained using bcftools [9] 'isec -n=2 -w 1' between the two VCF files. Finally, the population wide pseudo-testcross set was filtered for quality and missingness using vcftools ('--minDP 5, --minGQ 20, --max-missing 0.7'). The VCF was then parsed into a tab delimited file using GATK VariantsToTable and imported into R for further analysis. The phenotype data for each line were imported and lines were designated as resistant or susceptible as described above. A sample of 125 of the most CMD resistant and most susceptible (Resistant, both years' disease rating = 1; Susceptible, both years' disease rating >= 4) lines was randomly selected as the

Resistant and Susceptible Bulks, respectively, to perform the *in silico* bulk segregant analysis using the QTLseqr package[10]. For each SNP, the mean alternative allele ratio (SNP-index) for each bulk was calculated from all the individuals in the bulk and the difference in allele ratios was compared between the two bulks (ΔSNP-index). A 5Mb window tricube-smoothed ΔSNP-index was compared to the 95% confidence interval as in Takagi et al., 2014. SNPs with ΔSNP-index values surpassing the 95% confidence interval are significantly linked to the resistance phenotype.

Fine mapping using GBS and KASP markers: Field phenotyping and GBS markers are both susceptible to experimental error. In the field, susceptible plants may be scored as resistant if they do not experience sufficient disease pressure. Similarly, SNPs called from GBS data may be biased from low coverage or repetitive regions of the genome. A KASP marker-based assay with a second ~1000 individual population phenotyped by an extremely accurate VIGS based approach[11]. A window of 1.5 Mb bracketing the GBS defined QTL was targeted for analysis. Pseudo-testcross positions were then identified by aligning WGS reads from both NASE14 and two TME204-LCR lines to the TME204 hap1 assembly and examining the reads in the two parental lines and selecting heterozygous locations which have high complexity and minimum 30% GC content in the 100 bp surrounding the SNP. Primers were then designed by IDT using their PACE/KASP marker submission form (Supplementary Table 5). The standard KASP protocol was used to genotype every individual in the fine mapping population on a BioRad CFX384 using the Allelic Discrimination tab in the CFX software package.

The full $F_1$ fine-mapping population was phenotyped and after genotyping, recombinant lines were identified in the full population between markers M1, M2 and M6, M8 and these lines were further screened using the markers within that interval (M3, M5, M7). The original marker numbering scheme represents their order based on the AM560-2 ref 6.1 assembly, however the positions have been updated to reflect the more accurate positions in the TME204 hap1 assembly. The list of recombinants was narrowed to only those with phenotype-genotype mismatch and a minimal recombination site was identified as linked to the phenotype. To confirm these results, 5-7 replicates of each line were regenerated from tissue culture and re-phenotyped sing the above methods. The genotypes of the regenerated lines were also confirmed with all

KASP markers, and the recombinant lines and controls were sequenced using Illumina as above to verify the genotype and recombination locations.

Monte Carlo Sampling

After performing the SNP analysis, the number (n) of SNPs leading to an amino acid change were counted for the given scenario. Next, we randomly chose n bp positions throughout all 18 chromosomes of the TME204 genome, and marked them as being hypothetical SNPs. If at least one SNP was present within the defined locus (between marker M3 and M7), we identified this iteration of the experiment to have yielded a success otherwise, the round was counted as being unsuccessful. We repeated this experiment 100’000 times and the ratio of successes represents a rough estimate of the likelihood that an amino acid changing SNP is found by chance within a locus.

Virus Induced Gene Silencing

VIGS vector construction and plant inoculation: The VIGS-based screening method developed by Lentz *et al*. (2018)[12] was used to study the effects of the gene of interest on CMD resistance. Sequence from *MePOLD1* CDS (Manes.12G077400, 400 bp) was cloned into the multiple cloning site of the ACMV-based VIGS vector with *KpnI* and *SpeI*[12]. The DNA fragments were designed based on the TME204 reference genome (Qi et al., 2021) and the sequence was tested for specificity in 60444 (Kuon *et al*., 2019). Between fifteen to thirty plants were injected with *Agrobacterium tumefaciens* AGL1 containing VIGS constructs as described by Lentz *et al.* (2018)[12]. ACMV titre quantification: Total DNA was extracted from the top 1-2 leaves. Leaves were harvested at first signs of CMD symptoms and snap frozen in liquid nitrogen with the DNeasy Plant Mini Kit (QIAGEN, Germany). Quality was assessed by Nanodrop (Thermo Scientific, Wilmington, USA) and quantified with Qubit dsDNA BR Assay Kit (Thermo Fisher Scientific Inc, Massachusetts, USA). ACMV titre was quantified with qPCR using the LightCycler 480 System (Roche) with 15 ng of total DNA, 1 µM of primers, and Fast SYBR Green Master Mix (Applied Biosystems, Massachusetts, USA) in a final volume of 10 µL. ACMV DNA-A specific primers and the endogenous cassava *PP2A* gene (Manes.09G039900) was used as an internal control (Supplementary Table 4) and at least three technical replicates were included per sample. A Mann-Whitney U test was used to analyse the statistical significance. Primers are listed in Supplementary table 7.

Gene expression analysis: Total RNA was extracted from the top 1-2 leaves. Leaves were harvested at first signs of CMD symptoms and snap frozen in liquid nitrogen with the Spectrum Plant Total RNA Kit (Sigma-Aldrich, Merck Life Science, Germany) according to Protocol A. An On-column DNAase I Digestion (Sigma-Aldrich, Merck Life Science, Germany) was performed as manufacturer's instructions to remove residual genomic DNA. RNA quality was assessed with the Nanodrop system (Thermo Scientific, Wilmington, USA) and quantified with Qubit RNA BR Assay Kit (Thermo Fisher Scientific Inc, Massachusetts, USA). The samples were converted to cDNA using the RevertAid First Strand cDNA Synthesis Kit (Thermo Scientific, Wilmington, USA) according to the manufacturer's instructions. MePOLD1 (Manes.12G077400) relative expression was quantified with RT-qPCR in triplicates using the LightCycler 480 System (Roche) with 15 ng of cDNA, 1 µM of primers, and Fast SYBR Green Master Mix (Applied Biosystems, Massachusetts, USA) in a final volume of 10 µL. The comparative $C_T$ (threshold cycle) method (Livak and Schmittgen, 2001) was used to calculate relative transcript levels with Tubulin 1 β chain (*MeTUB1*, Manes.08G061700) as the reference gene. Primers are listed in Supplementary table 7.

7. Van der Auwera, G. A. & O'Connor, B. D. *Genomics in the Cloud: Using Docker, GATK, and WDL in Terra*. ("O'Reilly Media, Inc.," 2020).

**Supplementary Table 1: Genotypes of plants used for EWAS analysis.**

| Name | Genotype | ID | Phenotype | Reads No. | Coverage* | Conversion rate** |
|---|---|---|---|---|---|---|
| TME 7 WT | TME7 | cBS14p | Resistant to CMD | 846983521 | 114 | 99.40% |
| TME 7 WT | TME7 | cBS8p | | 859059800 | 115.62 | 99.42% |
| TME 7 WT | TME7 | cBS15p | | 844574034 | 113.67 | 99.40% |
| TME7 OES | TME7 | cBS10p | Susceptible to CMD | 853329597 | 114.85 | 99.41% |
| TME7 OES | TME7 | cBS11p | | 774679128 | 104.27 | 99.41% |
| TME7 OES | TME7 | cBS12p | | 775731790 | 104.41 | 99.49% |
| TME7 Organogenesis | TME7 | cBS13p | Resistant to CMD | 726885049 | 97.83 | 99.55% |
| TME7 Organogenesis | TME7 | cBS7p | | 832384407 | 112.03 | 99.52% |
| TME7 Organogenesis | TME7 | cBS9p | | 760693488 | 102.38 | 99.53% |
| TME7 Organogenesis | TME7 | cBS16p | Susceptible to CMD | 794495267 | 106.93 | 99.39% |
| TME7 Organogenesis | TME7 | cBS17p | | 754538912 | 101.56 | 99.41% |
| TME7 Organogenesis | TME7 | cBS18p | | 744368702 | 100.19 | 99.43% |
| TME204 WT | TME204 | TME204 | Resistant to CMD | 1606930158 | 223.43 | 99.70% |
| TME204 FEC | TME204 | FEC | Susceptible to CMD | 1659955923 | 233.43 | 99.76% |

WT: wildtype plant, OES: plant regenerated from organized embryogenic structure, Organogenesis: plant regenerated via caulogenesis, FEC: plant regenerated from friable embryogenic callus. * Estimated with genome size, ** Estimated with chloroplast

**Supplementary Table 2. Primers used**

| Name | Sequence (5' - 3') | Purpose |
|---|---|---|
| ACMV_F | GGTCCTGGATTGCAGAGGAAGATAGTGGG | Quantification of ACMV (DNA-A) genomic DNA by qPCR |
| ACMV_R | GGTACAACGTCATTGATGACGTCGATCCC | |
| MePOLD1_F | GCCAACTTGAGTTTGATTGCCTG | Quantification of MePOLD1 (Manes.12G077400) cDNA by qPCR |
| MePOLD1_R | GGATCATGGGTAGGCTC | |
| MePP2A_F | CGCTGTGGAAATATGGCATCA | Quantification of MePP2A (Manes.09G039900) genomic DNA by qPCR |
| MePP2A_R | CTGGCTCAAACTGCAGGATCAA | |
| MeTUB1_F | TGCCATGTTCCGTGGAAAGATG | Quantification of MeTub1 (Tubulin 1 β chain, Manes.08G061700) cDNA by qPCR |
| MeTUB1_R | CCCCTAGGTGGAATGTCACAGACAC | |
| TME204_F1 | TTATCTTCTGTGGCCCTTTTTC | Sanger analysis of MePOLD1 codon for amino acid 528 |
| TME204_R1 | TCAGGAGTTGAGAAAGTACC | |
| TME3_91_02324_F2 | GCTTATTATTCCTTTGTGCG | Sanger analysis of MePOLD1 codons for amino acids 680 and 685 |
| TME3_91_02324_R2 | CTTTCTATAAGCTTTGGCCT | |
| MePOLD1cds_F | TCACTCTCACTCTCACTCTCATTA | Amplify full length MePOLD1 cds |
| MePOLD1cds_R | CAGATTAGAAGTTCCACCTATCCA | |

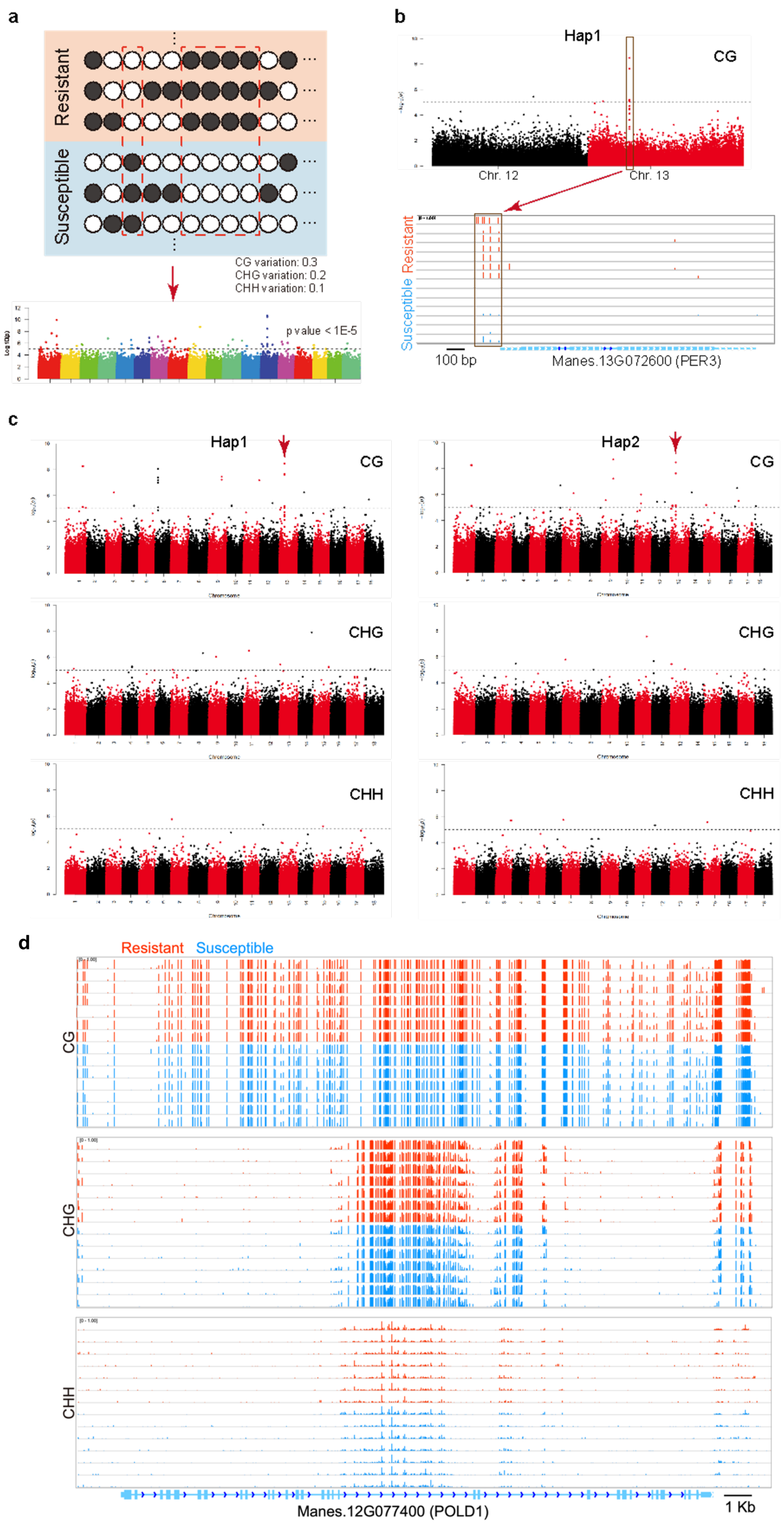


**Supplementary Fig. 1.** Full genome bisulfite sequencing of cassava varieties before and after tissue culture induced *de novo* morphogenesis. A) Analysis strategy cartoon. B) A single peak on chromosome 13 was identified as differentially methylated between resistant and susceptible lines. C) differential methylation across two haplotypes (Qi, W. et al 2021) for CG, CHG and CHH methylation. D) *Me POLD1* does not display differential methylation between resistant and susceptible cassava lines.

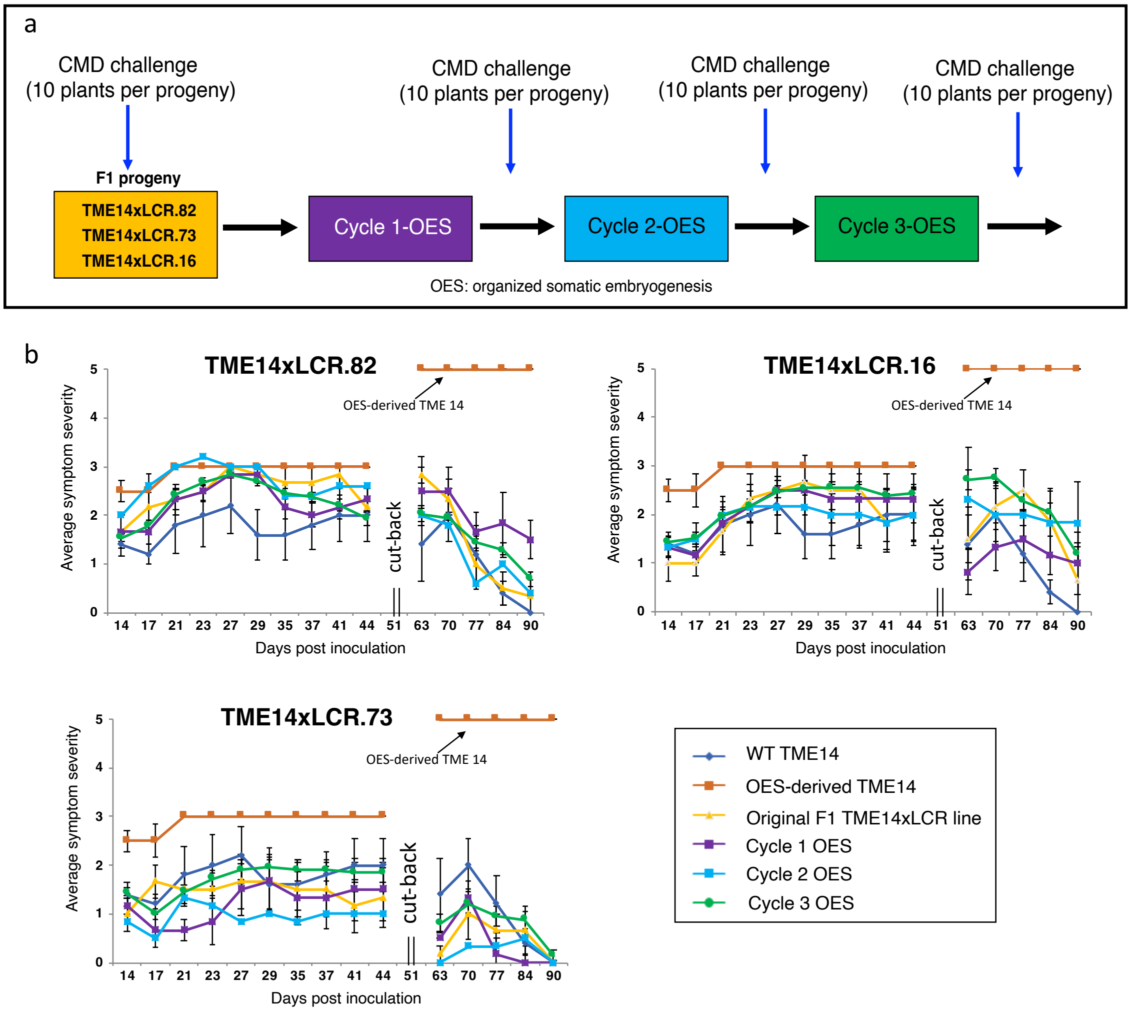


**Supplementary Fig. 2**. CMD2-mediated resistance remains stable in cassava progeny lines generated through sexual crosses. TME204-LCR lines were crossed with TME14 which carries functional CMD2-type resistance. a. Three progeny lines were passed through successive cycles of somatic embryogenesis. Plants were regenerated after each cycle and inoculated with the virulent EACMV isolate K201. b. CMD leaf symptoms were assessed visually on a 0-5 scale over 44 days, after which plants were ratooned and new growth scored until 90 days after inoculation. TME14 wildtype plants (blue, identical data shown in all three panels) displayed the recovery phenotype typical for this landrace, while OES-derived TME14 (LCR) lines passed through embryogenesis (orange, identical data shown in all three panels) became highly CMD susceptible, displaying the highest disease severity (5) and no recovery (arrowed). Plants regenerated from TME14xLCR F1 progenies remained resistant to CMD at a level not significantly different from the original F1 progeny line from which they were derived. OES: organized embryogenic structures

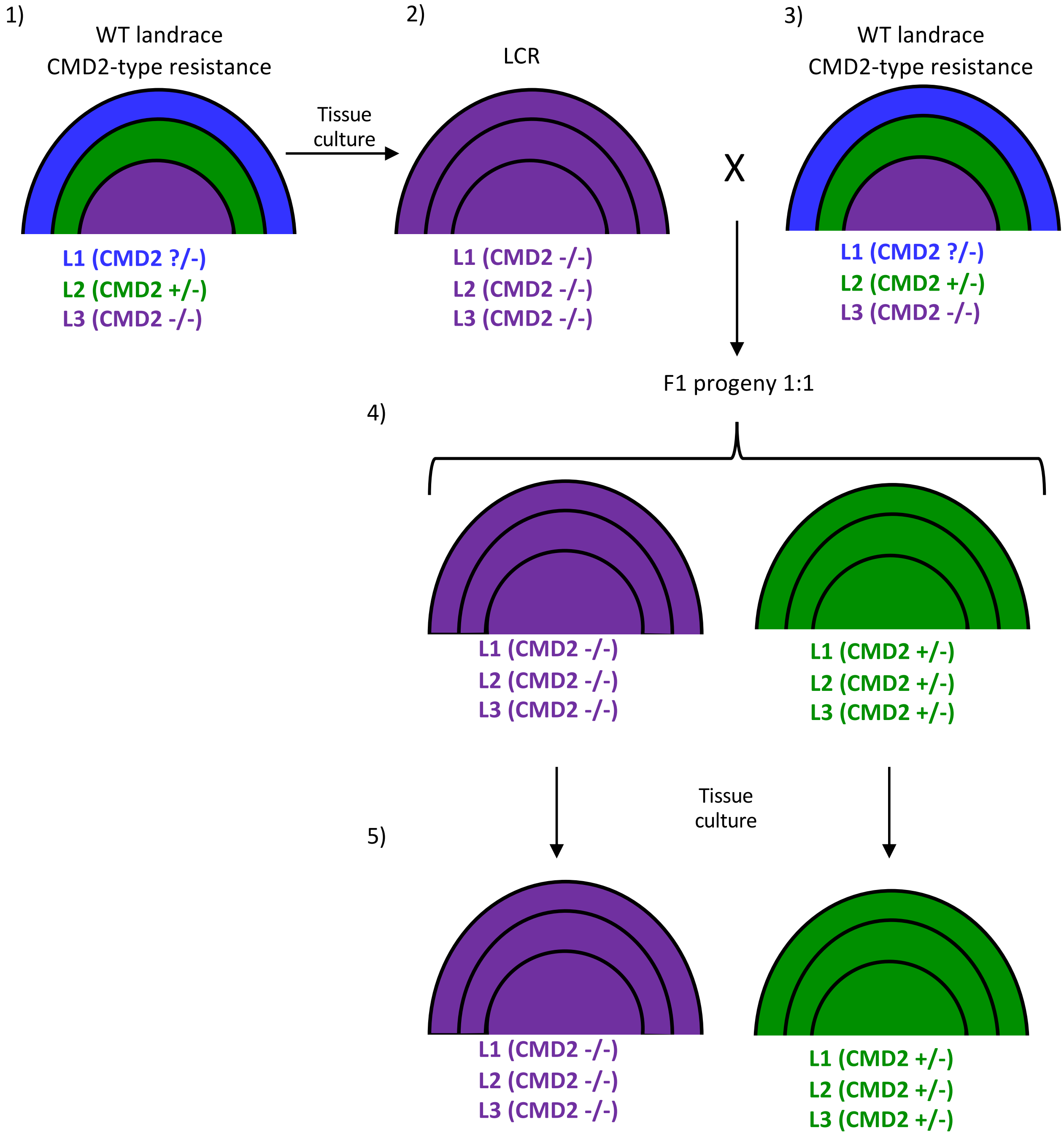


**Supplementary Fig. 3.** In the model cartooned below, L2 contains the resistant allele (green) while L3 does not (purple) (1). L3 gives rise to embryogenetic tissue (Loss of CMD2 Resistance - LCR) through *de novo* morphogenesis that is non chimeric and lacks the resistance allele (2). A similar scenario is possible if the L1 is -/- for CMD2 and gives rise to embryogenic tissue. When an LCR plant is crossed with a chimeric wildtype (WT) CMD2-type cassava variety, F1 progeny match the genotype of the parental L2 layer and therefore segregate resistance 1:1 and are not chimeric (4). Consequently, when resistant and susceptible F1 progeny are passaged through tissue culture induced *de novo* morphogenesis, the maintain their resistant or susceptible phenotypes, respectively.

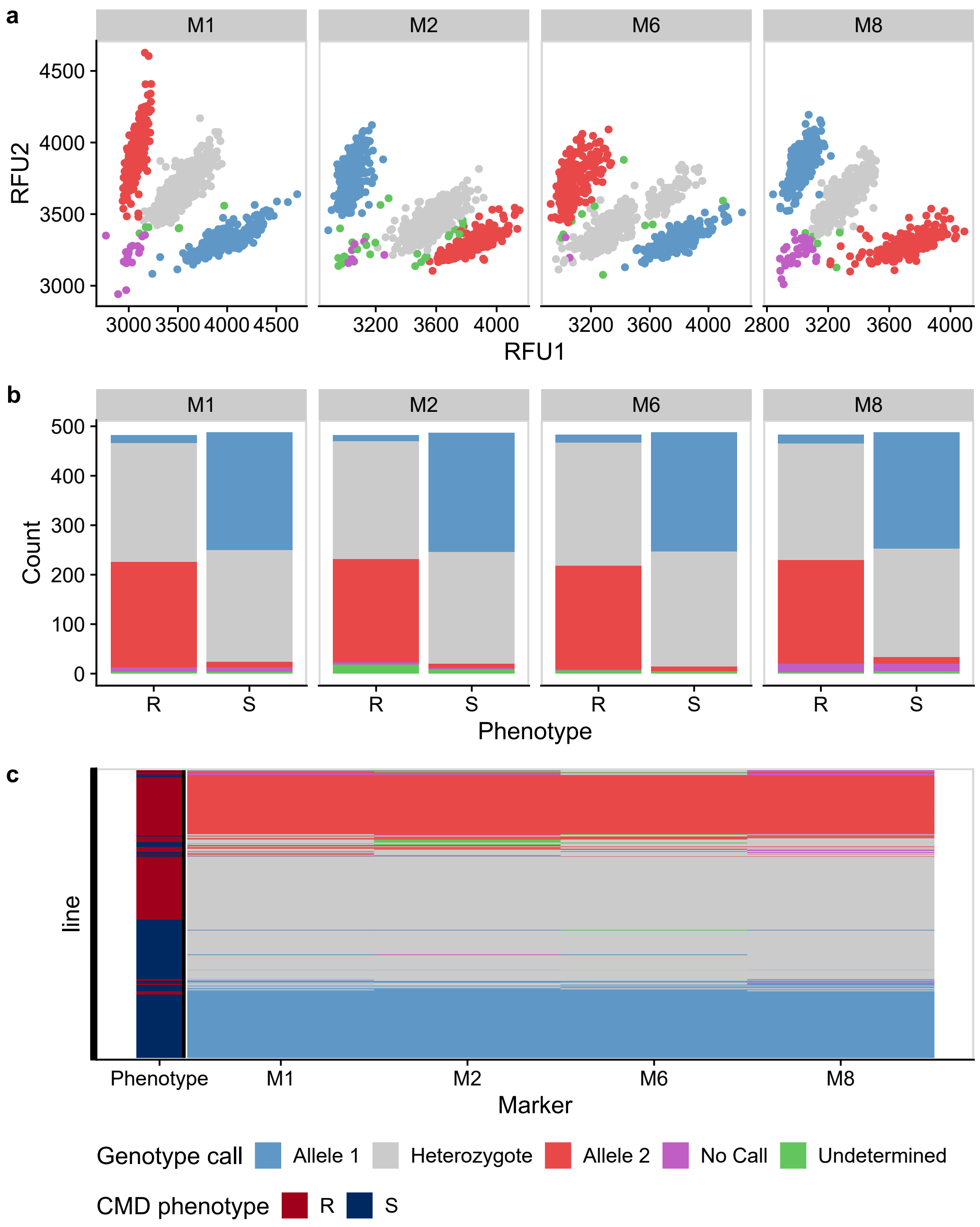


**Supplementary Fig. 4**. Genotyping using Kompetitive Allele Specific PCR (KASP) markers. a) The entire ~1000 F1 population was screened using 4 KASP markers, M1, M2, M6, and M8. Dot plot of the raw relative fluorescence units (RFU) for the two allele specific primers. Each point is an individual F1 progeny and the colours represent the allele call made for each marker. b) The distribution of genotype calls for each of the two phenotypic states (R - Resistant, S - Susceptible). c) A view of the all the calls of the entire ~1000 individual population. The markers on the x-axis are ordered by their genomic position allowing to visualize recombinants between the markers. The resistance phenotype is indicated on the left bar.

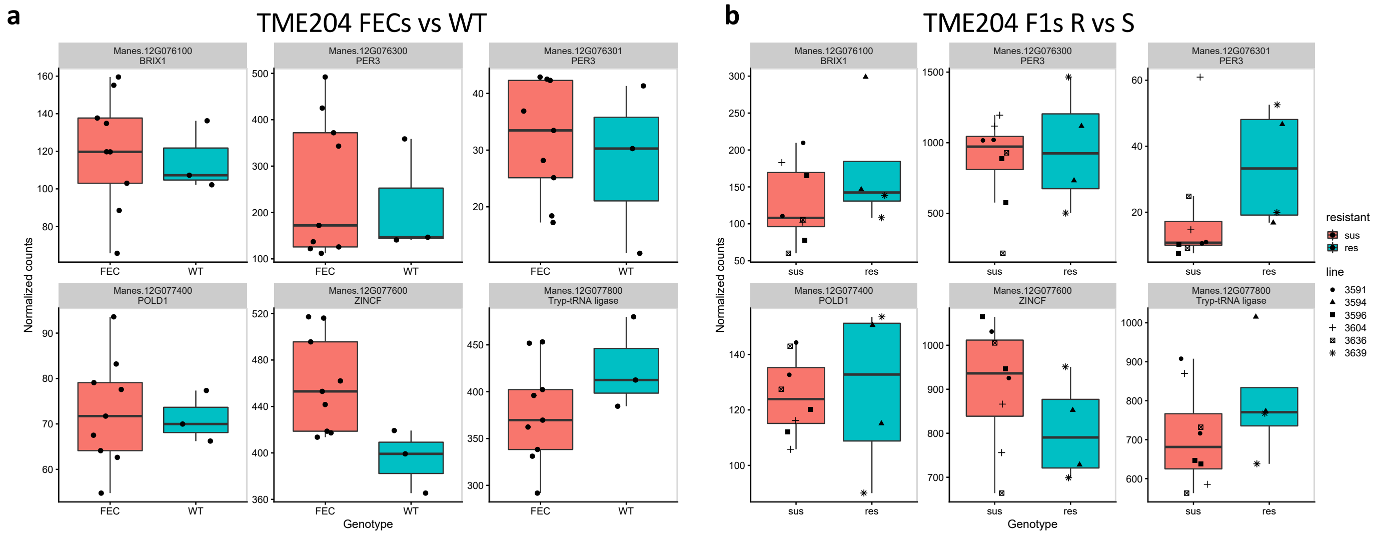


**Supplementary Fig. 5.** Gene expression analysis of genes within the fine-mapped CMD2 locus. (A) Resistant TME204 wildtype plants were compared to susceptible TME204-LCR plants regenerated through embryogenesis during production of friable embryonic callus (FEC). (B) Resistant and susceptible F1 plants derived from a TME204-WT self-cross were compared. Of the 8 genes defined within the 190Kb locus, only 6 are detected as expressed in our datasets. BRIX1 - RIBOSOME BIOGENESIS PROTEIN BRIX; PER3 – PEROXIDASE 3-RELATED; POLD1 - DNA POLYMERASE DELTA; ZINCF - ZINC FINGER, CCCH-type.

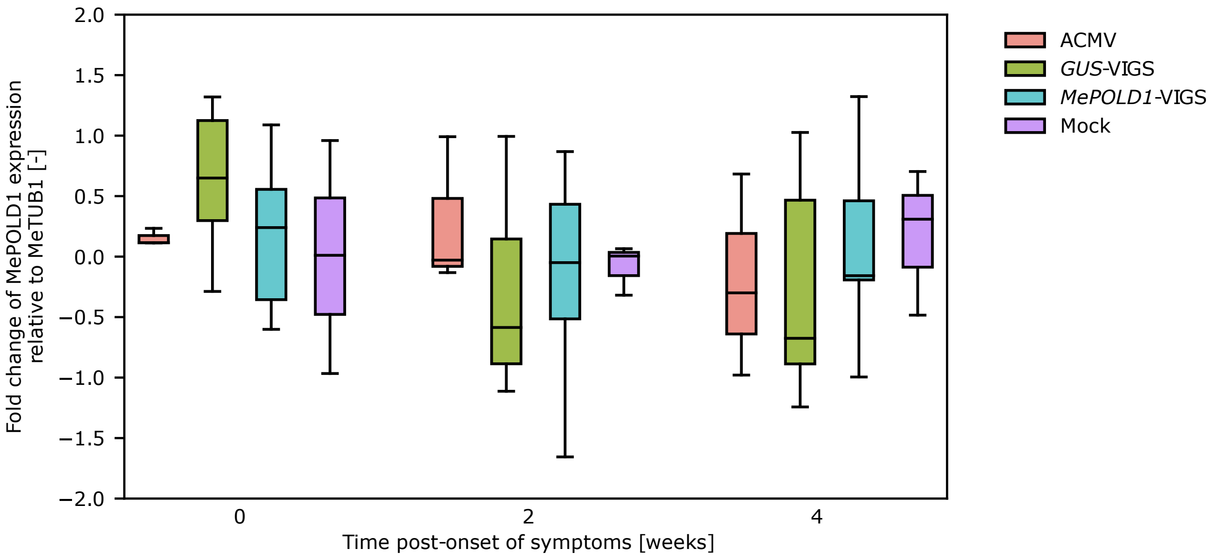


**Supplementary Fig. 6.** *MePOLD1* expression relative to expression of the gene encoding *Tubulin 1 β chain* (*MeTUB1*, Manes.08G061700) after ACMV-VIGS inoculation (non-modified ACMV, *GUS*-VIGS, *MePOLD1*-VIGS and mock) of CMD- susceptible cassava 60444. Week 0 is the first onset of symptoms of individual plants and week 2 is two weeks after that and so on. Number of biological replicates at Week 0, 2 and 4 respectively: ACMV (n = 3, 3, 3), *GUS*-VIGS (n = 10, 10, 8), *MePOLD1*-VIGS (n = 9, 10, 9), and mock (n = 3, 3, 3).

*MePOLD1*

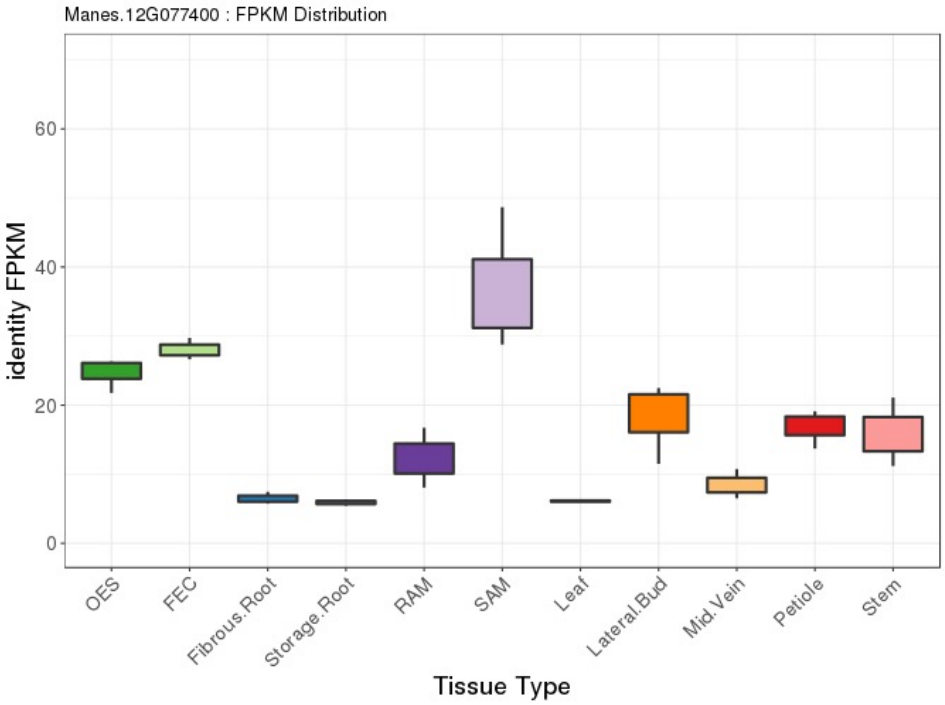


*MeTUB1*

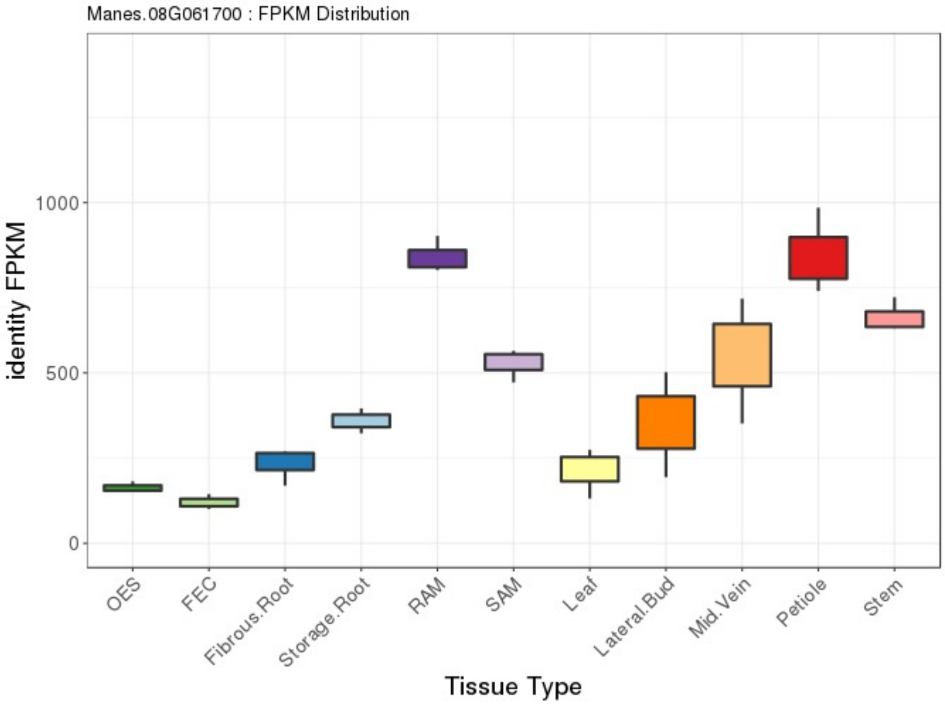


**Supplementary Fig. 7.** Gene expression across multiple tissue types. Data obtained from the Cassava Expression Atlas (Wilson et al. 2017).

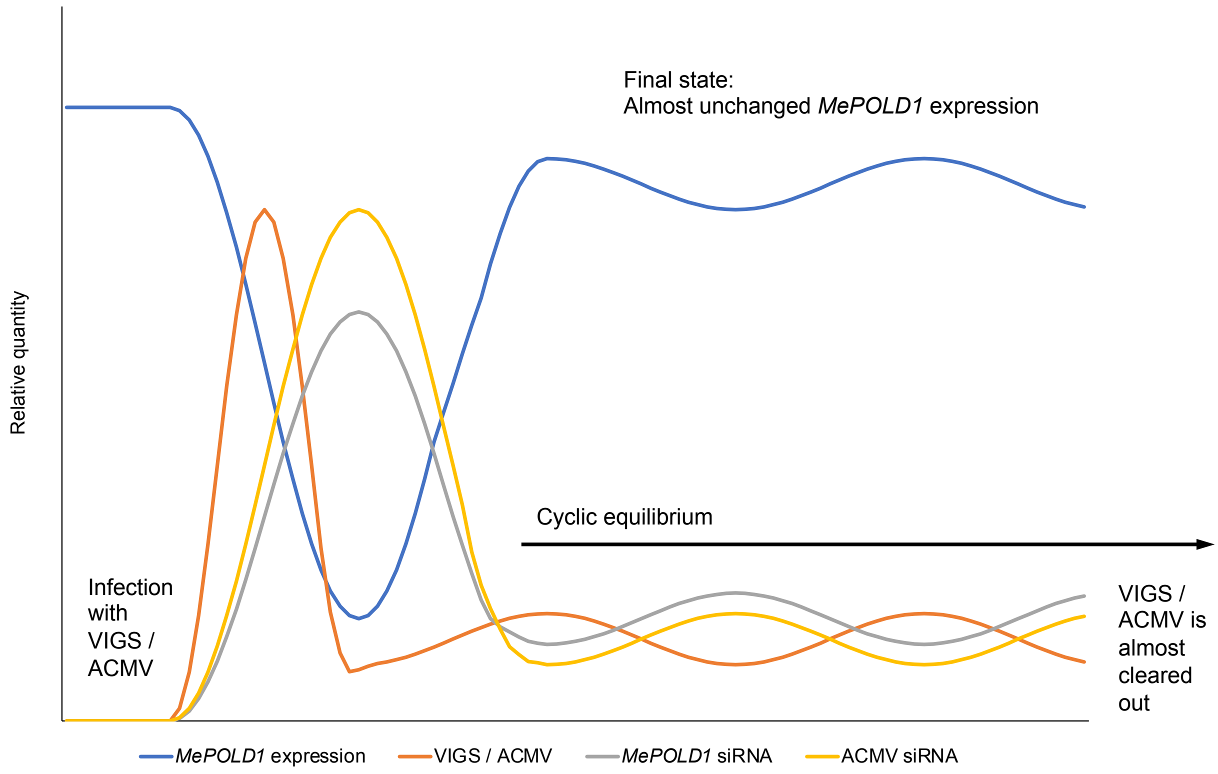


**Supplementary Fig. 8.** Hypothesis to explain the observed *MePOLD1* expression pattern after ACMV-VIGS inoculation and reduction of ACMV load in CMD-susceptible 60444**.** The blue line represents the hypothetical *MePOLD1* expression and it starts off under normal conditions. Once the plant has been inoculated with the *MePOLD1* VIGS construct which is a modified ACMV clone (represented by the orange line – VIGS/ACMV), the rise of VIGS/ACMV leads to the reduction (increase in *MePOLD1* siRNA represented by the grey line as well as siRNA of ACMV represented by the yellow line) of *MePOLD1* expression. Since ACMV needs MePOLD1 to replicate, as *MePOLD1* expression drops and ACMV siRNA increases, the quantity of VIGS/ACMV will also decrease. The reduction in VIGS/ACMV then leads to lower production of siRNA against *MePOLD1* thereby allowing the expression of *MePOLD1* to return to approximately normal quantities. Since *MePOLD1* level returns close to normal, VIGS/ACMV will also begin increase in quantity. However, since there are residual amounts of ACMV siRNA, VIGS/ACMV will never be able to establish itself thus leading to a cyclic equilibrium where VIGS/ACMV is maintained at a low quantity and *MePOLD1* expression remains almost unchanged.

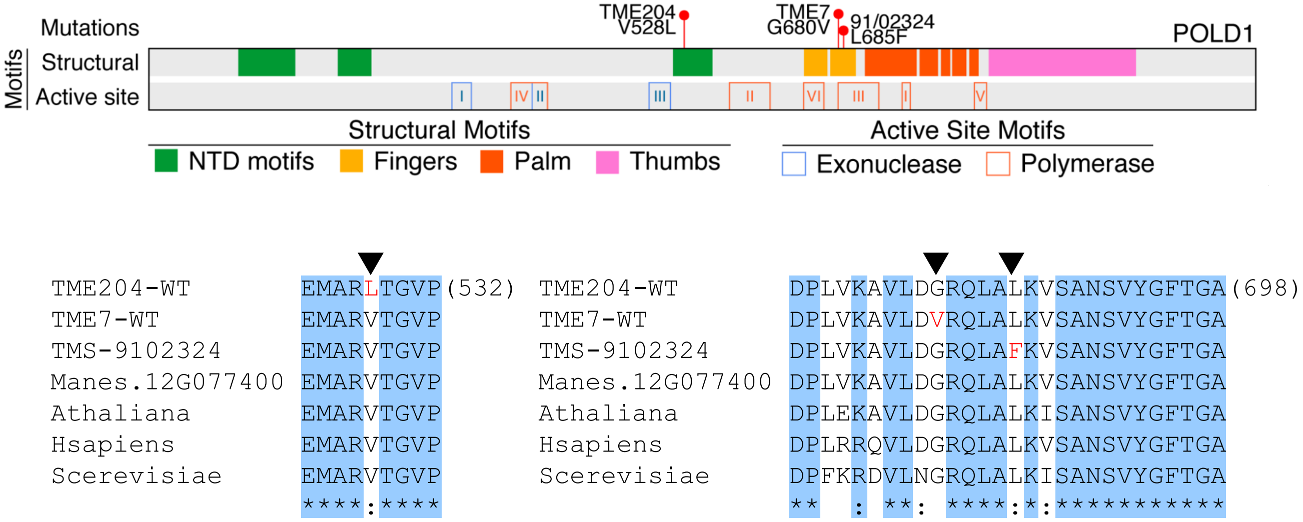


**Supplementary Fig. 9**. (Top) Schematic diagram of the POLD1 protein from cassava. Red lollipop flags indicate locations of resistance alleles. Active site motifs in the exonuclease and polymerase domains are indicated by the blue and orange outlined boxes, respectively. (Bottom) Protein alignment of POLD1 sequences. Sequences from three varieties containing a non-synonymous SNP are included. Affected amino acid is noted by an arrow head and the mutated residue in red; position of the last amino acid in alignment is indicated in paratheses. Manes.12G77400: *Manihot esculenta* AM560-2 v6.1; Athaliana: *Arabidopsis thaliana*; Hsapiens: *Homo sapiens*; Scerevisiae: *Saccharomyces cerevisiae*.

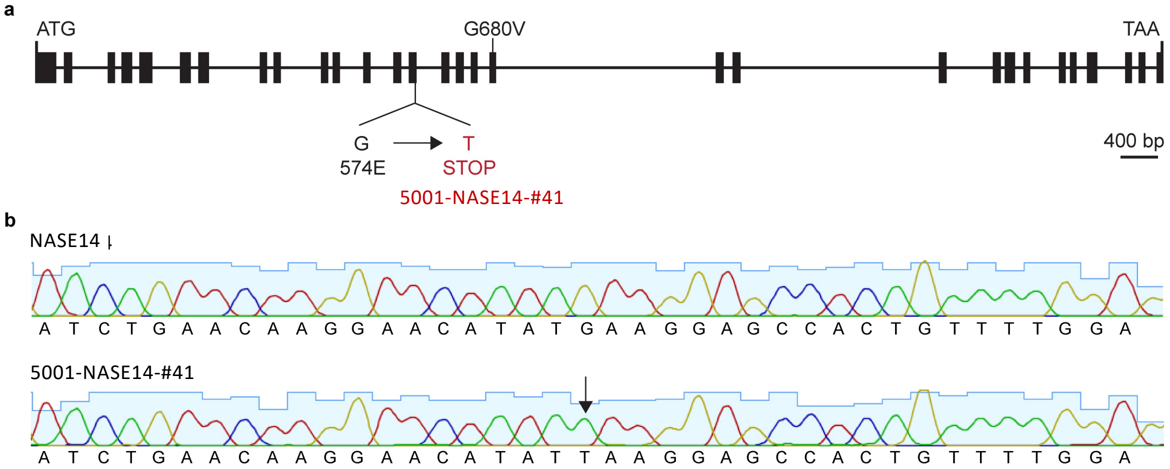


**Supplementary Fig. 10.** Premature stop codon within the resistant haplotype of *MePOLD1* in susceptible line 5001-NASE14-#41. 5001-NASE14-#41 is a transgenic line from resistant NASE14. **a**, Schematic diagrams show the gene structure of the resistant haplotype of *MePOLD1.* The exons are indicated as solid boxes, and the introns are indicated as lines. The mutation site in the resistant haplotype of *MePOLD1* in line 5001-NASE14-#41 is highlighted in red. **b**, Sanger sequencing analysis identified a mutation in the resistant haplotype of *MePOLD1* in line 5001-NASE14-#41. The full-length cDNA sequence was amplified by a pair of primers which specifically worked for resistant haplotype of *MePOLD1* in resistant NASE14 and its derived lines.
